# Toxin/Antitoxin Systems Induce Persistence and Work in Concert with Restriction/Modification Systems to Inhibit Phage

**DOI:** 10.1101/2023.02.25.529695

**Authors:** Laura Fernández-García, Sooyeon Song, Joy Kirigo, Michael E. Battisti, Daniel Huelgas-Méndez, Rodolfo García-Contreras, Maiken E. Petersen, María Tomás, Thomas K. Wood

## Abstract

Myriad bacterial anti-phage systems have been described and often the mechanism of programmed cell death is invoked for phage inhibition. However, there is little evidence of ‘suicide’ under physiological conditions for these systems. Instead of death to stop phage propagation, we show here that persister cells, i.e., transiently-tolerant, dormant, antibiotic-insensitive cells, are formed and survive using the *Escherichia coli* C496_10 tripartite toxin/antitoxin system MqsR/MqsA/MqsC to inhibit T2 phage. Specifically, MqsR/MqsA/MqsC inhibited T2 phage by one million-fold and reduced T2 titers by 500-fold. During T2 phage attack, in the presence of MqsR/MqsA/MqsC, evidence of persistence include the single-cell physiological change of reduced metabolism (via flow cytometry), increased spherical morphology (via transmission electron microscopy), and heterogeneous resuscitation. Critically, we found restriction-modification systems (primarily EcoK McrBC) work in concert with the toxin/antitoxin system to inactivate phage, likely while the cells are in the persister state. Phage attack also induces persistence in *Klebsiella* and *Pseudomonas* spp. Hence, phage attack invokes a stress response similar to antibiotics, starvation, and oxidation, which leads to persistence, and this dormant state likely allows restriction/modification systems to clear phage DNA.

## INTRODUCTION

Phage attack, resulting in up to 40% bacterial cell death in oceans (1), is the universal constraint on bacterial immortality; hence, bacteria have developed numerous phage-inhibition systems. For phage inhibition, there is little evidence that these systems cause cell death; instead, it is much more likely these systems invoke dormancy (2,3). Bacterial cell dormancy has several potential benefits: (i) allowing time for other host-encoded, phage-inhibition systems to function, (ii) slowing viral replication, and (iii) accommodating for spacer accumulation for CRISPR-Cas (2). In fact, the very existence of spacers in CRISPR-Cas type III and VI systems; i.e., those shown to induce dormancy by targeting host RNA, demonstrate cells must survive phage attack to obtain spacers. Also, growth-inhibited cells have more spacer acquisition (4). In contrast, programmed cell death during phage attack is paradoxical from the perspective of individual cells and has yet to be proven beneficial for kin.

Given that cells enter a state of dormancy known as persistence to survive numerous stresses (e.g., antibiotics (5), starvation (6), oxidation (7)) through guanosine pentaphosphate and tetraphosphate (p)ppGpp signaling (8), which leads to dimerization of their ribosomes to cease translation (9-11), we reasoned that phage attack is another form of extreme stress, so it should lead to persistence, in some cases. Also, phage-inhibition systems include toxin proteins of toxin/antitoxin systems, which are known to induce persistence by up to 14,000-fold (7,12,13). For example, after transcription shutoff by T4 phage activates the Hok/Sok toxin/antitoxin system, Hok damages the cell membrane to cease cell metabolism to halt phage propagation, in the first link of a toxin/antitoxin system to phage inhibition (1996) (14). Similarly, phage attack activates various toxins to reduce cell metabolism including those of the CBASS (phospholipases, nucleases, and pore-forming proteins (15)), RADAR (RdrB converts ATP to ITP (16)), and Thoeris (ThsA degrades NAD^+^ (17)) phage inhibition systems.

Therefore, based on these two insights (extreme stress produces persisters and phage-inhibition systems utilize toxins to reduce metabolism), we investigated whether persister cells are formed as a result of inducing the *bona fide Escherichia* spp. phage inhibition system, the tripartite toxin/antitoxin/chaperone system *E. coli* C496_10 MqsR/MqsA/MqsC (18,19), which utilizes RNase MqsR (20,21) upon phage attack. We confirmed that the MqsR/MqsA/MqsC, produced from its natural promoter, via the same system used to identify it, dramatically inhibits T2 phage (19), and discovered this system converts *E. coli* into the persister state to inhibit phage. Similar results of phage inducing persistence were found with *Klebsiella pneumoniae* and *Pseudomonas aeruginosa*. In addition, we discovered that restriction/modification systems assist the MqsR/MqsA/MqsC toxin/antitoxin system to inhibit T2 phage. To our knowledge, this is the is the first connection of phage inhibition with persistence, and the first non-CRISPR-Cas system to be linked to restriction/modification systems.

## MATERIALS AND METHODS

### Bacteria and growth conditions

Bacteria and plasmids are shown in **Table S1**, and cells were cultured at 37°C in LB supplemented with 30 µg/ml of chloramphenicol (to retain plasmids, LB Cm30). The complete MqsR/MqsA/MqsC plasmid-based system (pCV1-*mqsRAC* and pCV1 empty plasmid control) was verified via Plasmidsaurus sequencing.

### Persister formation assay and phage titers

Single colonies were cultured overnight in LB-chloramphenicol, then diluted 100X and grown to a turbidity of 0.5 at 600 nm. T2 phage was then added (multiplicity of infection ∼ 0.1) for one hour, then the cells were washed twice with phosphate buffered saline (PBS, 8 g of NaCl, 0.2 g of KCl, 1.15 g of Na_2_HPO_4_, and 0.2 g of KH_2_PO_4_ in 1000 mL of dH_2_O) and cultured for three hours with LB-ampicillin (100 µg/mL, 10 minimum inhibitory concentration (MIC)) or LB-ciprofloxacin (5 µg/mL, 10 MIC) to kill non-persister cells (11). Mitomycin C was added in LB (10 µg/mL, 5 MIC) to kill persister cells (22). Cells were washed twice with PBS and enumerated on LB-Cm plates to measure persistence. To determine phage titers, after 30 minutes of phage contact, supernatants were diluted with phage buffer (0.1 M NaCl, 10 mM MgSO_4_ and 20 mM Tris-HCl pH 7.5), and plaques were enumerated on top agar, TA, (1% tryptone, 0.5% NaCl, 1.5% agar lower layer and 0.4% agar top layer) double layer plates.

### Persister resuscitation assay

Single-cell microscopy of resuscitating persister cells was performed as described previously (23) using LB agarose gel pads that were observed up to 3 h via a light microscope (Zeiss Axioscope.A1, bl_ph channel at 1000 ms exposure). The persister cells were generated by the addition of T2 phage at 0.1 MOI for 1 h. The microscope was maintained at 37°C by placing it in a vinyl glove box (Coy Labs), which was warmed by an anaerobic chamber heater (Coy Labs, 8535-025).

### Metabolic activity via flow cytometry

To ascertain metabolic activity, *E. coli* MG1655/pCV1-*mqsRAC* was grown to a turbidity of 0.5 at 600 nm and T2 phage was added (MOI 0.1) for 1 h. Cells were washed, resuspended in PBS, stained for metabolic activity (BacLight™ RedoxSensor™ Green Vitality Kit, Thermo Fisher Scientific Inc., Waltham, MA, USA), which measures the cellular redox state. The fluorescence signal of 100,000 cells was analyzed by flow cytometry (Beckman Coulter LSRFortessa) using the FL1 (530 ± 30 nm).

### Transmission electron microscopy

To confirm persister cell formation, *E. coli* MG1655/pCV1-*mqsRAC* was grown to a turbidity of 0.5 at 600 nm, and T2 phage was added (MOI 0.1) for 1 h. Cells were centrifuged and fixed with 2% glutaraldehyde in 0.1 M sodium cacodylate buffer followed by secondary fixation with 1% osmium tetroxide. Samples were then stained with 2% uranyl acetate, dehydrated, and embedded in eponate. After curing, blocks were sliced with an ultramicrotome, and 70 nm thin slices were collected on grids and post-stained with uranyl acetate and lead citrate.

### qRT-PCR

To quantify transcription after T2 phage attack (15 min at MOI 0.1) in the presence of MqsR/MqsA/MqsC, RNA was isolated by cooling rapidly using ethanol/dry ice in the presence of RNA Later (Sigma) using the High Pure RNA Isolation Kit (Roche). The following qRT-PCR thermocycling protocol was used with the iTaq^TM^ universal SYBR^®^ Green One-Step kit (Bio-Rad): 95 °C for 5 min; 40 cycles of 95 °C for 15 s, 60°C for 1 min for two replicate reactions for each sample/primer pair. The annealing temperature was 60°C for all primers. Primers are shown in **Table S2**.

### Efficiency of plating and efficiency of center of infection assays

The efficiency of plating assay (24) was performed for *E. coli* by serially diluting phage T2 (dilutions from 10^-1^ to 10^-7^) onto double agar plates that were inoculated with 100 μL of overnight cultures and dried for a minimum 15 minutes. Diluted phage (5 μL) was added to soft agar plates, the plates were incubated overnight, and the number of plaque forming units (PFU)/mL was determined. The efficiency of the center of infection (ECOI) assay (25) was used to determine the ability of T2 phage to infect different strains (i.e., a pre-adsorbed productive infection assay) and performed by diluting overnight bacterial cultures 1:100 into 15 mL of fresh buffered LB Cm30 and incubating to a turbidity at 600 nm of ̴ 0.5, then T2 phage (MOI ∼0.1) was added to the cultures and incubated for 8 minutes (adsorption time). The cultures were washed twice with PBS (centrifuging at 5000 rpm for 10 min) to remove free phages. Then the cultures were diluted in phage buffer (dilutions 1 to 5), and 5 μL drops were added to soft agar plates containing MG1655. Plates were placed at 37 °C for overnight incubation and PFU/mL determined.

### Klebsiella pneumoniae and Pseudomonas aeruginosa persister assays after phage attack

*K. pneumoniae* strain obtain from blood infection (26) was infected with DNA lytic phage vB_KpnP-VAC25 (27) at MOI 0.01, washed with PBS, and contacted with 10 MIC colistin (10 µg/mL) or 5 MIC mitomycin C (10 µg/mL). *P. aeruginosa* PAO1 from the Gloria Soberón collection at UNAM was infected with phage PaMx12 (DNA lytic phage, sequence https://www.ncbi.nlm.nih.gov/nuccore/386649691) at MOI 1 and surviving colonies in the center of the plaques were isolated after 20 h and tested for phage sensitivity to determine if the cells were resistant or persistent to phage. To confirm persistence, cells were contacted with phage followed by lethal (10 MIC, gentamycin at 10 µg/mL for *P. aeruginosa*).

## RESULTS

Given the extensive experience with *E. coli* persister cells (7,9,22,28-33) and given our discovery of the MqsR/MqsA toxin/antitoxin system (20,21), we chose a phage exclusion system of *Escherichia* spp. based on MqsR/MqsA to investigate the hypothesis that persister cells are formed upon phage attack. We also used the same expression system that was used to demonstrate phage inhibition (via a plaquing assay) and used T2 phage since it was inhibited best by the MqsR/MqsA/MqsC system (19) and produced MqsR/MqsA/MqsC in a background that lacks this system (K-12).

### MqsR/MqsA/MqsC inhibits T2 phage

As published previously (19), T2 phage was efficiently inhibited by the MqsR/MqsA/MqsC TA system in our hands; i.e., MqsR/MqsA/MqsC increased cell viability by 10^6^-fold compared to an empty plasmid control (**Fig. 1**, **Table S3**). Corroborating the phage inhibition, the number of T2 phage decreased 500-fold in the presence of MqsR/MqsA/MqsC: 6 ± 2 x 10^5^ pfu/mL were obtained for MqsR/MqsA/MqsC whereas 2.6 ± 0.5 x 10^8^ pfu/mL were obtained for the empty plasmid when T2 was added initially at 7.14 x 10^6^ pfu (MOI of 0.1). Therefore, MqsR/MqsA/MqsC actively inhibits T2 phage under these conditions.

**Fig. 1.**
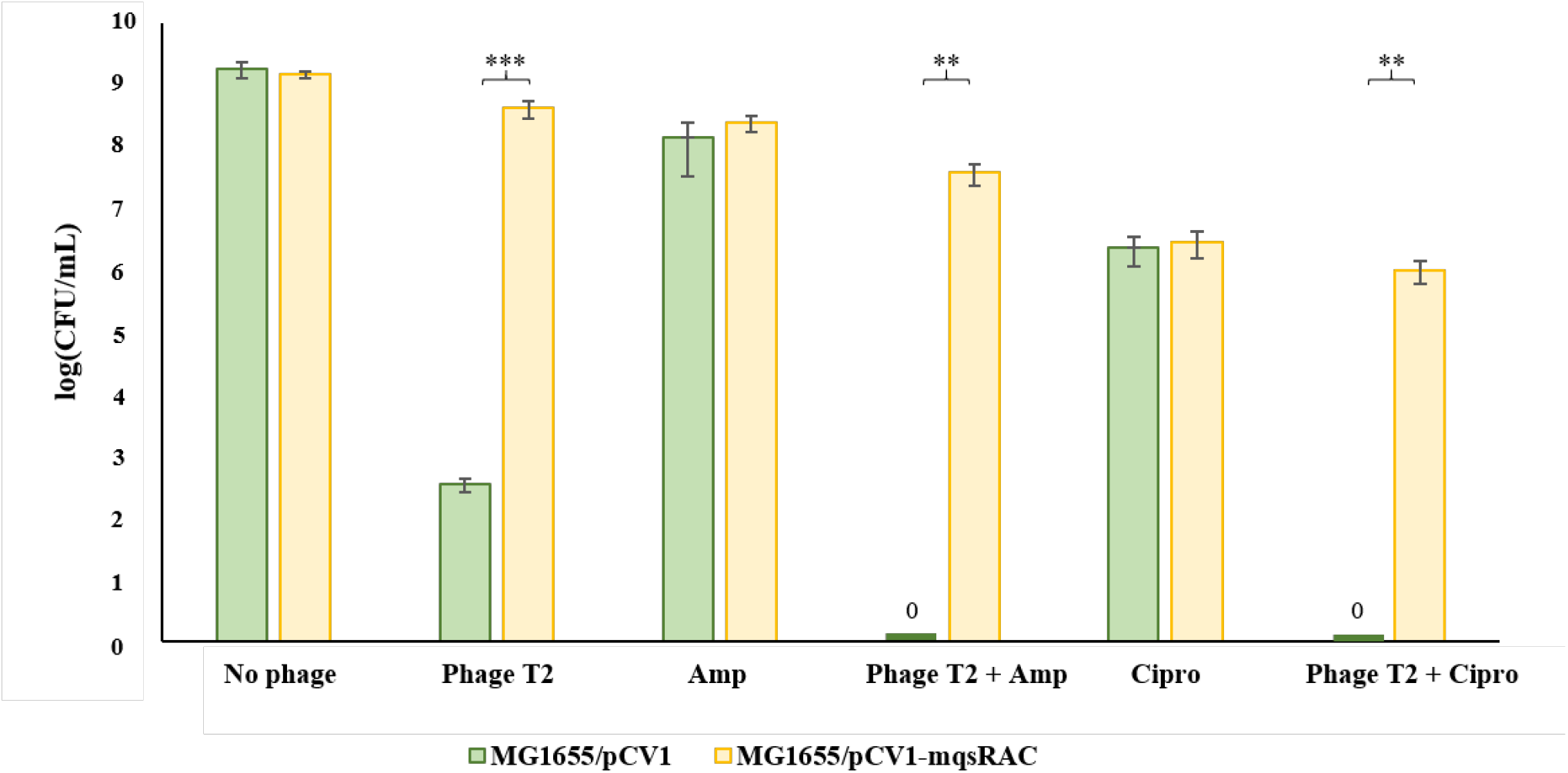
MqsR/MqsA/MqsC inhibits T2 phage by forming persister cells. Wildtype *E. coli* MG1655 cells producing MqsR/MqsA/MqsC from its native promoter (and the empty plasmid negative control) were contacted first with T2 phage at 0.1 MOI for 1 h followed by treatment with ampicillin (‘Amp’, 100 µg/mL, 10 MIC) or ciprofloxacin (‘Cipro’, 5 µg/mL, 10 MIC) for 3 h. Results from four independent cultures each. *** p-value < 0.005, **p-value < 0.01 (statistics via GradPath). One average deviation shown, and raw data are shown in **Table S3**.

To provide additional evidence of phage defense by the MqsR/MqsA/MqsC TA system, we performed both an efficiency of plating (EOP) and an efficiency of the center of infection (ECOI) assay. We found there was 2,100 to 5,000-fold reduction in the EOP with phage T2 when the cells produced MqsR/MqsA/MqsC (**Fig. 2A, Table S4A**). Consistently, we found the MqsR/MqsA/MqsC TA system reduces the ECOI by 5- to 14-fold (**Fig. 2B, Table S4B**). Hence, the MqsR/MqsA/MqsC TA system is a *bona fide* phage defense system.

**Fig. 2.**
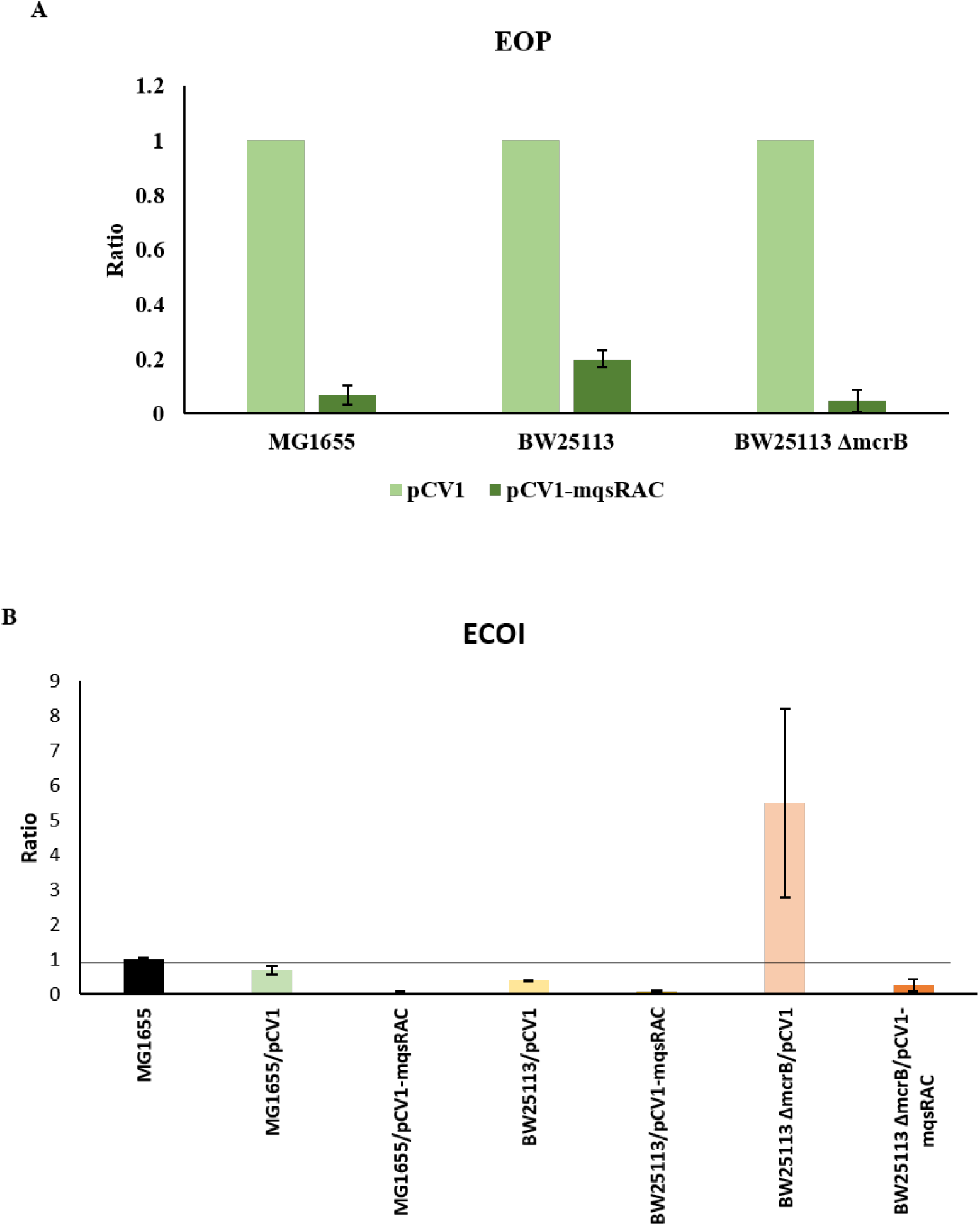
Efficiency of plating and efficiency (EOP) of the center of infection assay (ECOI). (**A**) EOP assay with light green columns showing results with empty plasmid pCV1 (‘no MqsRAC’), and dark green columns showing results with MqsR/MqsA/MqsC (‘MqsRAC’). (**B**) ECOI assay with the strains used in this manuscript. The efficiencies are relative to the strain MG1655. One average deviation shown, and raw data are shown in **Table S4**.

### Cells surviving T2 phage attack with MqsR/MqsA/MqsC are persisters

Critically, treatment with T2 phage (0.1 MOI) for one hour followed by treatment with 10X the minimum inhibitory concentration (MIC) of ampicillin induced persistence in 8% of the cells producing MqsR/MqsA/MqsC (**Fig. 1**, **Table S3**); i.e., these cells survived treatment of lethal ampicillin concentrations. In contrast, cells bearing the empty plasmid were completely eradicated by ampicillin after exposure to T2 phage indicating no cells entered the persister state without MqsR/MqsA/MqsC (**Fig. 1**). Normally, persister cells are formed at less than 1% of the population (34,35), so the number of persisters created by MqsR/MqsA/MqsC during T2 phage attack is striking (∼10%). Similar results were seen with a different class of antibiotic, ciprofloxacin (10X MIC), in that 0.3% cells with the MqsR/MqsA/MqsC phage inhibition system were persisters after T2 attack (**Table S3**), whereas no cells survived T2 and ciprofloxacin in the absence of MqsR/MqsA/MqsC (**Table S3**). Moreover, the increase in persistence by MqsR/MqsA/MqsC was not due to a difference in growth since *mqsRAC^+^* had virtually no impact on the specific growth rate in the absence of phage (**Fig. S1**, µ = 1.7 ± 0.1/h, µ = 1.62 ± 0.06/h, and µ = 1.74 ± 0.09/h for MG1655, MG1655/pCV1, and MG1655/pCV1-mqsRAC, respectively).

Since persister cells are metabolically inactive (5,31,36), we reasoned that if they are formed during T2 phage attack, they must resuscitate once the phage is removed and nutrients are supplied (23). Moreover, heterogeneity in persister cell resuscitation is a consistent, distinguishing feature of these cells (23,37-40). Therefore, we investigated the resuscitation of single cells on LB gel pads after 1 h of T2 attack (0.1 MOI) and found significant heterogeneity in waking with four phenotypes including 41% with immediate waking (less than 30 min), 33% waking with a lag (30 to 180 min), 11% elongating, and 15% waking then dying (data from representative sample) (**Fig. 3**, **Table S5**). In comparison, 97 ± 3% of the sample of exponentially-growing cells divide uniformly without elongation, lags, or death, forming microcolonies by 3 h (**Fig. S2**). Hence, during T2 attack, cells become persistent based on their antibiotic tolerance to two different classes of antibiotics and based on their heterogeneity in resuscitation.

**Fig. 3.**
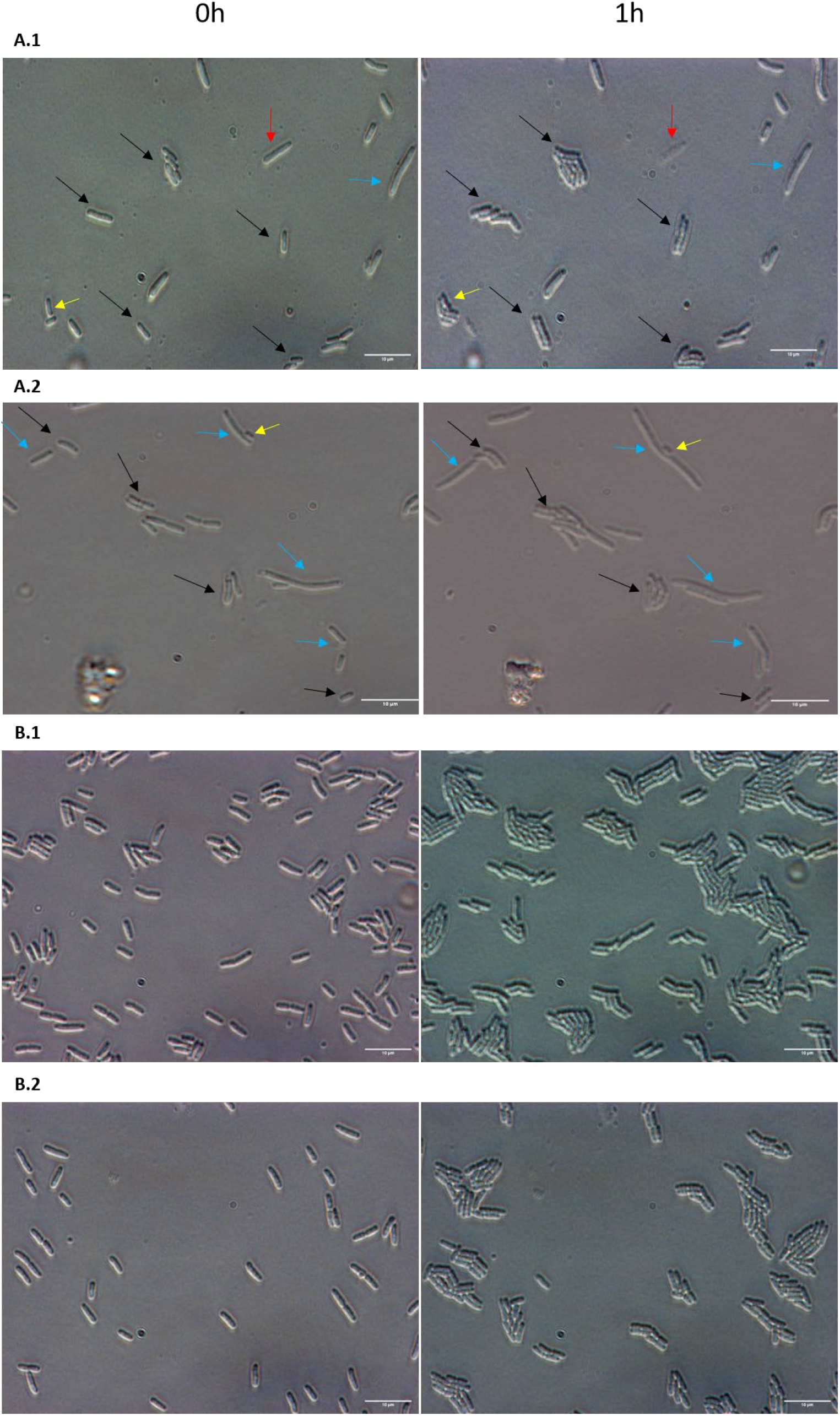
Heterogenous single-cell resuscitation after phage attack. Representative images (from five independent cultures) of the resuscitation of *E. coli* persister cells with MqsR/MqsA/MqsC (**A**) and division of exponential cells (**B**) after 1 h as determined with light microscopy (Zeiss Axio Scope.A1) using LB agarose gel pads (representative images from 0 to 3 h are shown in **Fig. S3**). The persister cells were generated by the addition of T2 phage at 0.1 MOI for 1 hour. Cells with the empty plasmid (i.e., no MqsR/MqsA/MqsC) are not shown due to the cellular debris that stems from complete eradication by T2 phage. Black arrows indicate cells with immediate waking (within 30 min), yellow arrows indicate cells with delayed waking (waking between 30 to 180 min), red arrows indicate cells that wake then die (lyse within 3 h), and blue arrows indicate cells that elongate initially. Black arrows were not added to exponential cultures pictures because all of them divide. Data for percentages are shown in **Table S5**.

Further evidence of persistence after T2 phage attack includes metabolic staining with flow cytometry, which revealed that after 1 h of T2 attack, *E. coli* cells with MqsR/MqsA/MqsC had significantly reduced metabolism compared to exponentially-growing cells (**Fig. S3**). This population with reduced metabolism has a broad range of metabolic activity since ∼25% of the cells remain viable after T2 attack for MG1655/pCV1-*mqsRAC*) but only ∼10% are dormant persisters (**Table S3**). In addition, transmission electron micrographs after T2 phage attack show the same features of persister cells seen previously after lethal antibiotic treatment and after starvation: dense cytosol, intact membranes, and spheroid shape (6) (**Fig. S4**).

### Persistence cell formation is rapid upon phage attack

To investigate how rapidly persister cells form upon phage attack, we investigated induction of the three loci required to inactivate ribosomes: *raiA, rmf,* and *hpf* (9) after 15 min of phage addition. Using qRT-PCR, we found no induction of these loci for the cells producing the MqsR/MqsA/MqsC phage inhibition systems (**Fig. 4**, **Table S6**), which indicates the cell probably uses existing proteins to become dormant rapidly, rather than relying on expression of ribosome inactivation genes once phage attacks. However, for cells that lack this phage inhibition system, *raiA* was induced 4-fold at 15 min after phage attack, which indicates these cells are stressed as they unsuccessfully try to thwart T2 phage. Bacteriophage T2 takes 60 minutes for its burst size, with a latent period of around 20 min and a rise period of 10 min (41).

**Fig. 4.**
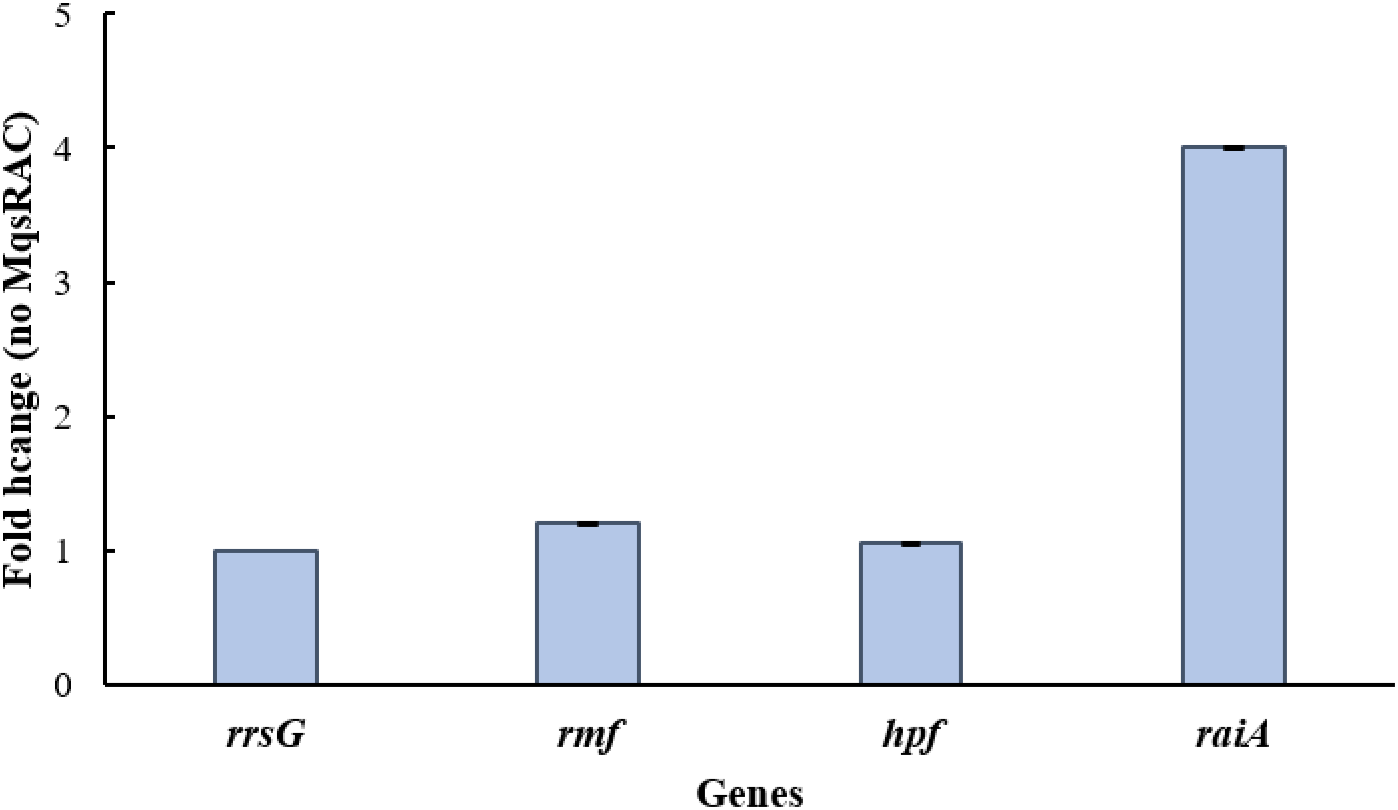
Phage attack does not induce ribosome inactivation proteins. Fold changes for wild-type *E. coli* with the empty plasmid pCV1 (‘no MqsRAC’) relative to the wild-type producing MqsR/MqsA/MqsC from pCV1-*mqsRAC*) after 15 min of T2 attack as determined by qRT-PCR. One average deviation shown.

### Restriction/modification assists MqsR/MqsA/MqsC during persistence

Since cells with MqsR/MqsA/MqsC weather T2 attack by rapidly becoming persistent and resuscitate, we hypothesized that another system works while the cells are dormant to clear T2 phage. Therefore, we investigated whether cells that lack the restriction systems/DNA processing enzymes encoded by *mrr, mcrB,* and *recD* are susceptible to T2 attack and found that inactivation of the McrBC system was most important for defending against T2 since its inactivation reduces cell viability by 10,000-fold (**Table S7A**). Critically, we found that the MqsR/MqsA/MqsC works with the McrBC restriction/modification system to defend against T2 phage since cells deficient in McrBC had 38-fold less viability, 10-fold less persister cells formed in the presence of MqsR/MqsA/MqsC (**Fig. 5, Table S7B**), and a 4-fold increase in PFU (**Table S7C**). Moreover, in the absence of MqsR/MqsA/MqsC, there was a 6-fold reduction cell viability after T2 phage attack when the McrBC restriction/modification systems was inactivated (**Fig. 5**, **Table S7B**). Furthermore, in the presence of MqsR/MqsA/MqsC, inactivating McrBC increases the efficiency of plating of T2 phage by 100-fold and increases the efficiency of the center of infection by 3-fold (**Table S4**). Therefore, MqsR/MqsA/MqsC works in concert with restriction/modification systems to inhibit T2 phage.

**Fig. 5.**
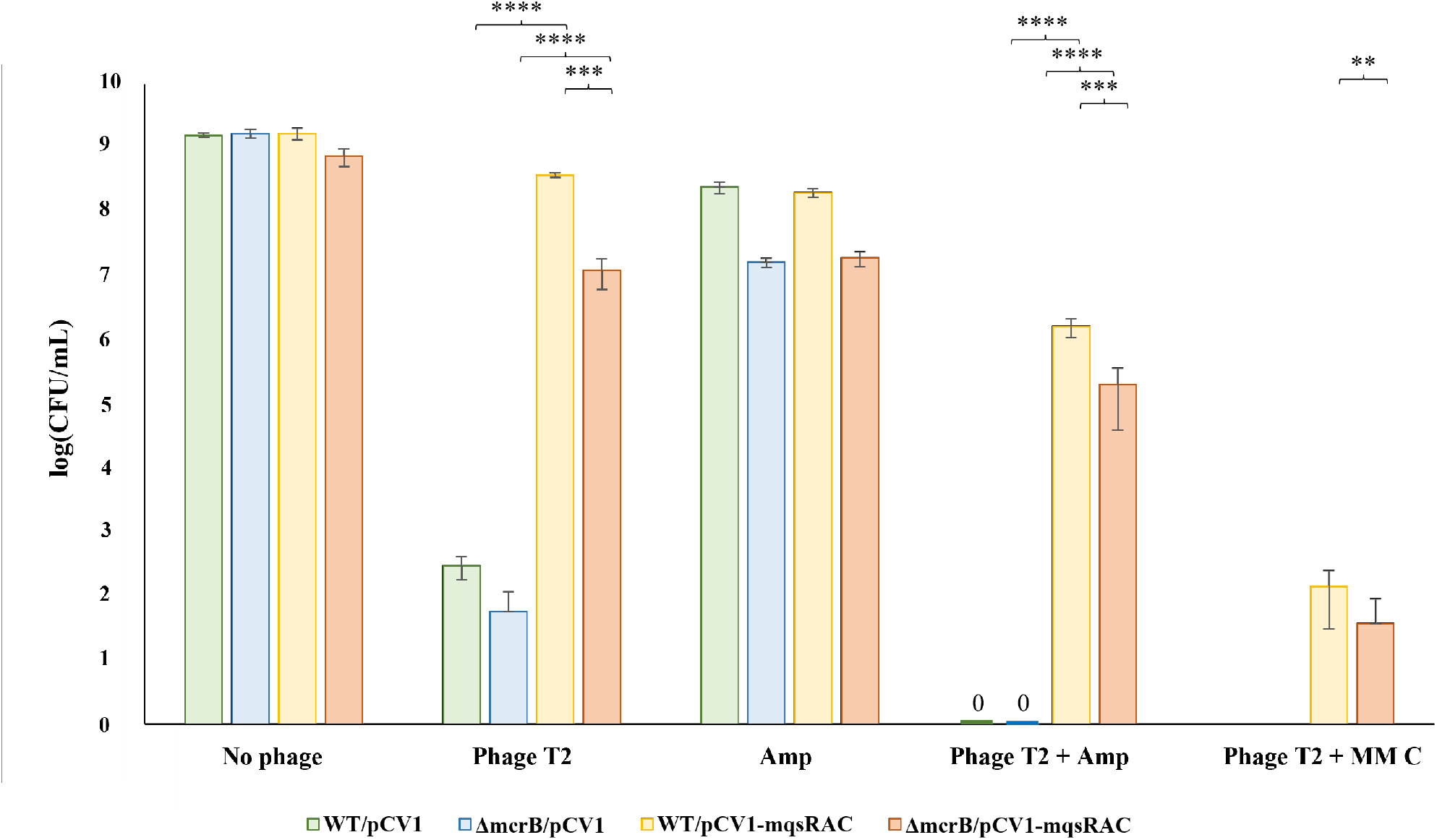
MqsR/MqsA/MqsC inhibits T2 phage through cooperation with restriction/modification systems. Wildtype *E. coli* and Δ*mcrB* cells producing MqsR/MqsA/MqsC from its native promoter (and the empty plasmid negative control) were contacted first with T2 phage at 0.1 MOI for 1 h followed by treatment with ampicillin (‘Amp’, 100 µg/mL, 10 MIC) or mitomycin C (‘MM C’, 10 µg/mL, 5 MIC) for 3 h. Results from four independent cultures each. **** p-value < 0.0001, *** p-value < 0.005, **p-value < 0.01 (statistics via GradPath). One average deviation shown.

### Mitomycin C kills *E. coli* phage-induced persister cells

Since phages are considered important for augmenting antibiotics to control infections (42), and since *E. coli* persister cells are formed rapidly during T2 phage attack, we investigated whether compounds that kill persister cells while they are dormant may be used to kill the remaining persisters. Approved by the FDA for cancer treatment, mitomycin C kills bacterial persister cells by passively diffusing into dormant cells and cross-linking their DNA (22) and has been shown to eradicate the pathogens *E. coli* O157:H7, *Staphylococcus aureus, Pseudomonas aeruginosa*, and *Acinetobacter baumannii* (22,43). We found that adding mitomycin C (10 µg/mL, 5 MIC) to MG1655/pCV1-*mqsRAC* for 3 h after 1 h of T2 treatment (MOI 0.1) led to a 100,000-fold reduction in viable cells (**Fig. 5)**. Similar results were obtained when McrBC was inactivated (**Fig. 5**). Hence, mitomycin C may be used to kill effectively the persister cells formed upon T2 phage attack.

### *K. pneumoniae* and *P. aeruginosa* form persister cells after phage attack

To explore whether persister cells are formed after phage attack in non-*E. coli* strains, we investigated whether phages induce persistence in *K. pneumoniae* and *P. aeruginosa.* For *K.* pneumoniae, we found that after treatment of 10^8^ cells/mL with VAC25 phage (0.01 MOI) for one hour, of the remaining viable cells after phage attack (10^4^ cells/mL), 11 ± 4% of the cells were persistent as shown by survival with 10X MIC of colistin for 3 h (**Fig. 6)**. In comparison, starting with the same initial cell density (10^4^ cells/mL) but omitting phage, 0 ± 0% the cells survived 3 h 10X MIC of colistin treatment (**Fig. 6)**. Furthermore, similar to *E. coli,* we found that mitomycin C eradicated the *K. pneumoniae* persister cells that were formed after VAC25 attack.

**Fig. 6.**
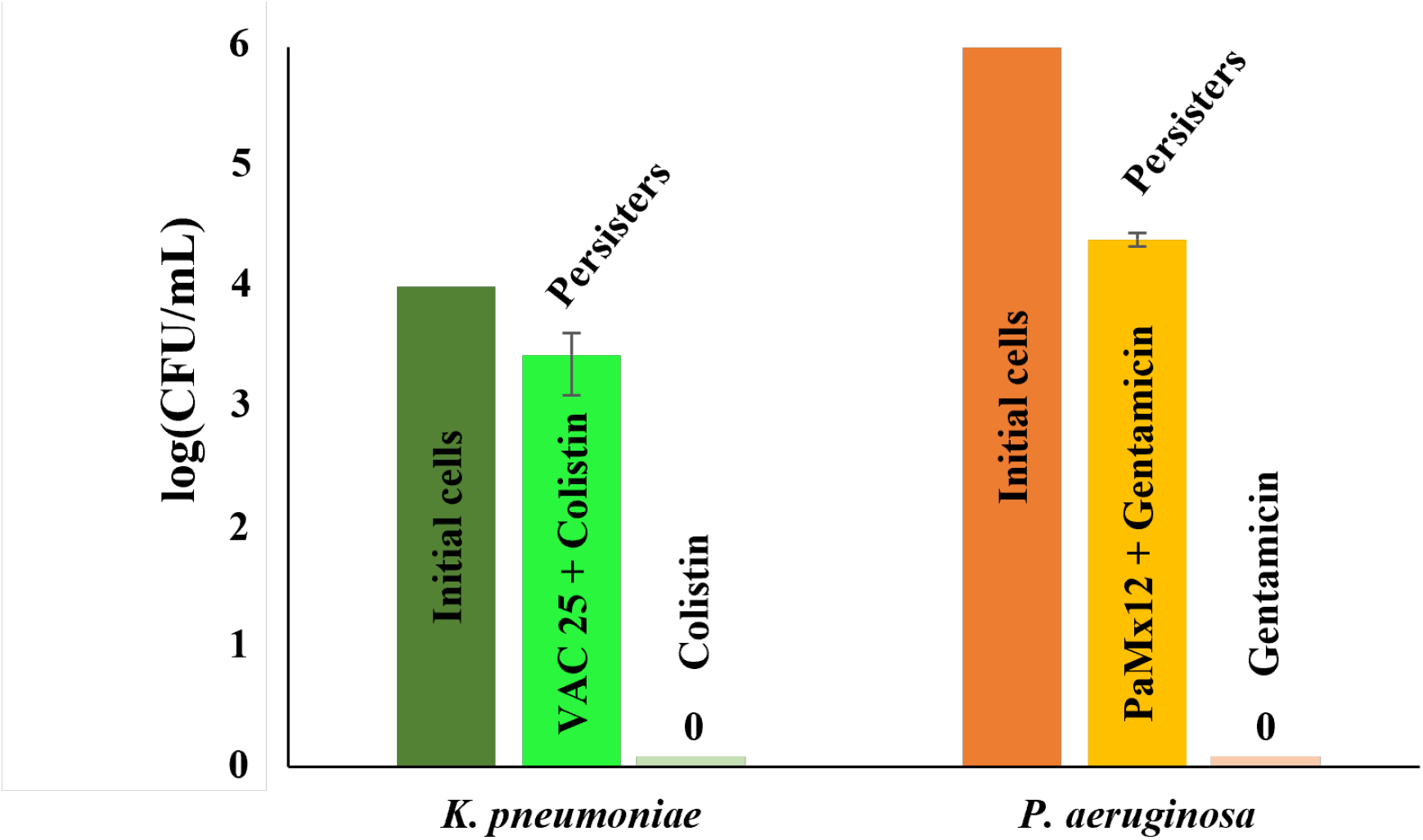
*K. pneumoniae* and *P. aeruginosa* form persister cells upon phage attack. First bar indicates initial cell density after phage attack (10^8^ *K. pneumoniae* cells/mL treated with 0.01 MOI VAC25 phage for one h or 10^8^ *P. aeruginosa* cells/mL treated with 1 MOI PaMx12 phage for one hour), second bar indicates phage and antibiotic treatment (0.1 MOI VAC25 phage for one h followed by 10X MIC of colistin for 3 h for *K. pneumoniae* or 1 MOI PaMx12 phage for one hour followed by 10X MIC of gentamicin for 3 h), and third bar indicates antibiotic treatment alone (10X MIC of colistin for 3 h added to 10^4^ cells/mL for *K. pneumoniae* or 10X MIC of gentamicin for 3 h added to 10^6^ cells/mL for *P. aeruginosa*). One average deviation shown.

For *P. aeruginosa*, we found that after treatment of 10^8^ cells/mL with PaMx12 phage (1 MOI) for one hour, of the remaining viable cells after phage attack (10^6^ cells/mL), 0.5 ± 0.14% cells were persistent as shown by survival with 10X MIC of gentamicin for 3 h. In comparison, starting with the same initial cell density (10^6^ cells/mL) but omitting phage, 0 ± 0% the cells survived 3 h 10X MIC of gentamicin. In addition, we found that 40% of the cells from the middle of plaques formed in soft agar are persistent, whereas 60% are resistant; i.e., 60% of the cells from the plaques developed mutations that reduced plaque formation in a subsequent assay with phage PaMx12. Hence, these results suggest the formation of persister cells during phage attack may be a general physiological response.

## DISCUSSION

Our results demonstrate the tripartite toxin/antitoxin MqsR/MqsA/MqsC phage inhibition system forms persister cells upon T2 phage attack. Moreover, this persistence allows at least some of the cells to survive phage attack by providing time for phage DNA degradation by restriction/modification systems. Hence, our results suggest toxin/antitoxin systems act first to reduce metabolism, then restriction/modification systems are utilized to undermine phage attack. Similarly, CRISPR-Cas has been shown to act first, followed by restriction/modification systems in *Listeria* spp. (44). Our results showing synergy between toxin/antitoxin and restriction/modification systems are significant since both of these phage defense systems are found in nearly all bacteria (unlike many other phage inhibition systems such as CRISPR-Cas, CBASS and retrons that are found in 39%, 13% and 11% of procaryote genomes, respectively (45)), which suggests this interaction is widespread and may be dominant for phage inhibition. Our results also provide the first reproducible link between toxin/antitoxin systems and persister cells under near-physiological conditions (native promoter and copy number 5 vector used (19,46)).

These results also have implications for other phage-inhibition systems in that, rather than causing cell death (47-50), these anti-phage systems probably induce dormancy; i.e., persistence, which allows the cell to thwart the phage in the same manner as we studied here. Along with the formation of persister cells, some cells indubitably succumb to phage attack; however, our results indicate the primary purpose of the anti-phage systems is not what has been termed ‘abortive infection’ in which the phage inhibition system kills the cells (19,47), but instead, these systems protect the cells and allow them to survive phage attack. Similar results showing dormancy have been seen with type VI CRISPR-Cas and restriction/modification systems (44).

Corroborating our findings, many phage-inhibition systems reduce energy metabolites like ATP (RADAR (16) and Detocs (51)) and NAD^+^ (Thoeris (17)), and the reduction of energy metabolites like ATP consistently induces persistence (11,52). Furthermore, reduction of ATP by toxin/antitoxin systems leads to persistence (53).

Central to persister cell formation is accumulation of (p)ppGpp (9-11); hence, it is expected that if cells prevent phage propagation by becoming persisters, they will do so via an increase in (p)ppGpp. Therefore, to counter this bacterial defense, if our hypothesis/data are correct in this report, phages should try to reduce (p)ppGpp levels. Critically, this has been recently shown with *Pectobacterium* spp. phage PcCB7V, which encodes the nucleotide pyrophosphatase MazG that is enriched against TIR-STING anti-phage systems and reduces (p)ppGpp (54). In addition, these authors demonstrated that (p)ppGpp is increased during phage attack (54).

Our model (**Fig. 7**) is that by invoking persistence (i.e., dormancy) through phage attack, back-up phage inhibition systems like restriction modification systems can degrade the phage genome since restriction enzymes do not require any resources from the cell for activity; i.e., Gibbs free energy is less than zero for phage DNA degradation due to the increase in entropy, so degradation of the phage genome can occur while the cells sleep. This general concept of allowing back-up DNA-targeting enzymes to function while the cell is dormant has been proposed for the type III/VI CRISPR-Cas systems (2,55) and demonstrated while this work was being prepared for type VI CRISPR-Cas in *Listeria* spp. (44); however, the benefit of dormancy has not been suggested for the plethora of recently-discovered, non-CRSPR-Cas phage inhibition systems. Here we demonstrate that (i) dormancy in the form of persistence is a key aspect of phage inhibition of non-CRISPR-Cas systems and that (ii) restriction/modification is utilized during the dormant state.

**Fig. 7.**
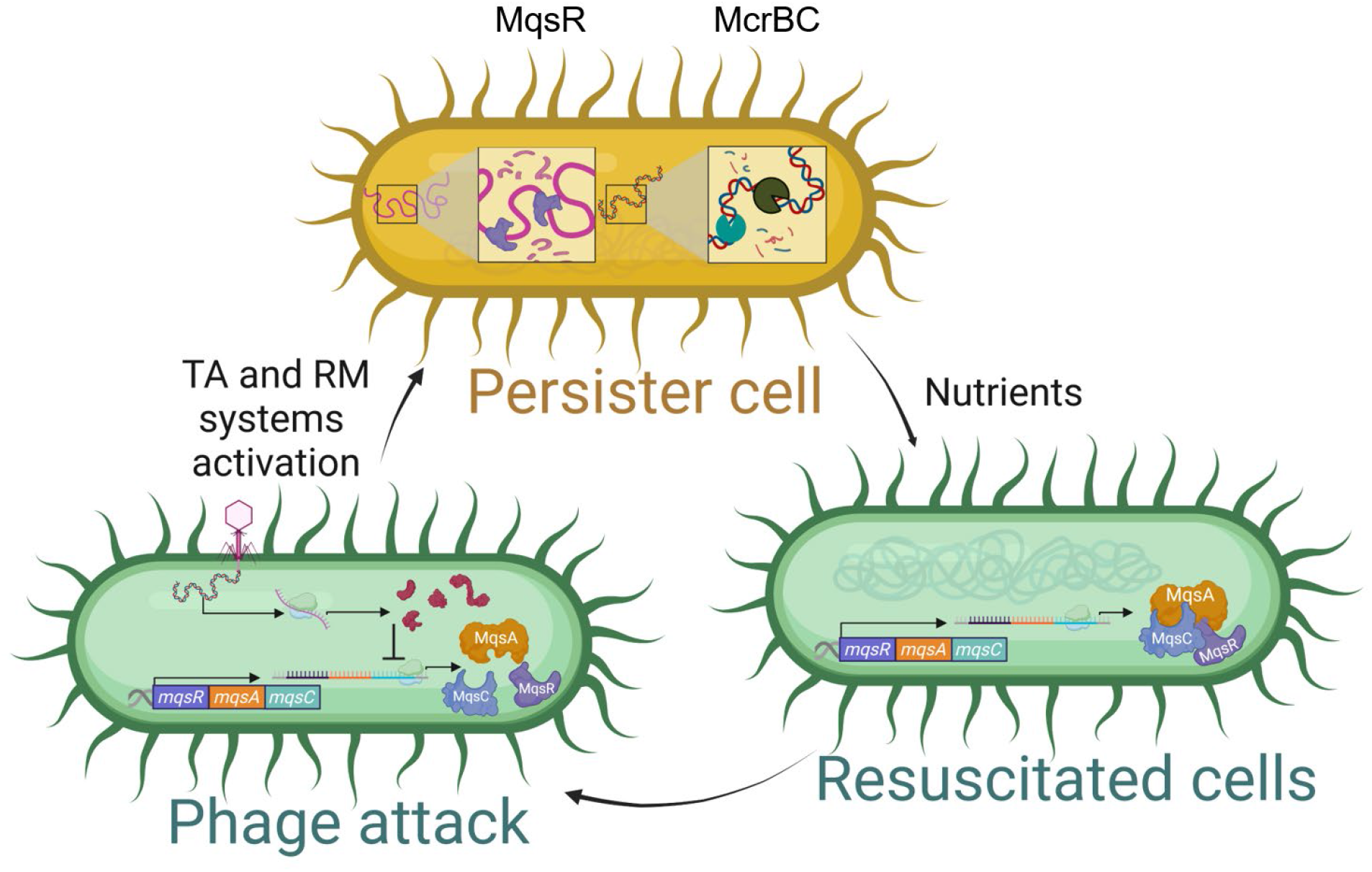
Schematic illustrating phage inhibition systems invoke dormancy. Our results indicate that after phage attack, bacteria become persister cells (i.e., dormant), through RNase decay via toxin MqsR of the MqsR/MqsA/MqsC tripartite toxin/antitoxin system. During this period of inactivity, restriction/modification systems, which do not require cell resources, degrade phage DNA. Upon favorable conditions (nutrients provided), the persister cells resuscitate, having survived the phage attack.

Since phages rarely eradicate populations, a logical extension of our results is that for Bacteria and Archaea, phage attack should lead to a small population of surviving cells in the persister state, since only a small population of cells survive other stresses like starvation and antibiotics (34,35). Our results showing phages induce persistence in both *K. pneumoniae* and *P. aeruginosa* lend credence to this generalization. Hence, for medical applications, a combination treatment will likely be required that recognizes that persister cells will develop during phage therapy. Therefore, compounds such as mitomycin C, which is capable of killing a wide range of cells in the persister state (22), will likely be required, as we demonstrate (**Fig. 5, Table S7B**).

Since the cells are becoming dormant (i.e., persisters) rather than dying, our results also suggest that after phage attack, the cells which survive phage, resuscitate upon detecting nutrients, as we have shown using single cell microscopy for persister cells (9,10,30), by using chemotaxis proteins and sugar transport membrane proteins, by reducing cAMP and ppGpp internal signals, and by increasing HflX activity to convert dimerized ribosomes (100S ribosomes) into active 70S ribosomes.

Finally, our results show that, like myriad other stresses such as starvation (6), oxidation (7), and antibiotic challenge (11), bacteria form persister cells to survive the extreme stress of phage attack. Hence, the formation of persister cells is a consistent and conserved means to weather stress.

## DATA AVAILABILITY

The data underlying this article are available in the article and in its online supplementary material.

## ACKNOWLEDGEMENTS

We are grateful for the MqsR/MqsA/MqsC plasmids provided by Prof. Michael Laub and for assistance from Missy Hazen of the Microscopy and Cytometry team from The Huck Institutes of the Life Sciences at PSU.

## Supporting Information

**Table S1.**
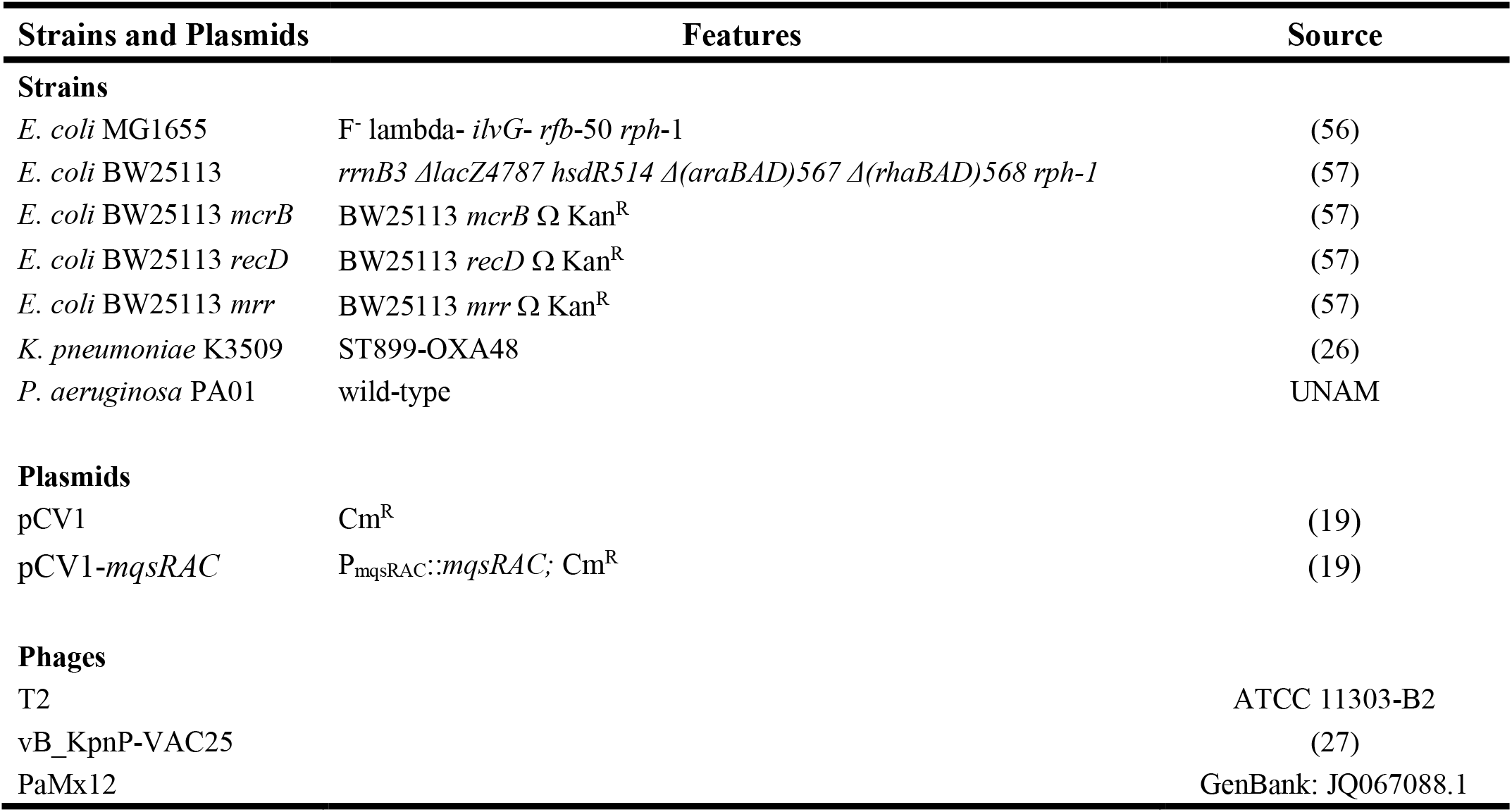
Bacterial strains and plasmids utilized. Cm^R^ is chloramphenicol resistance, and Kan^R^ is kanamycin resistance.

**Table S2.**
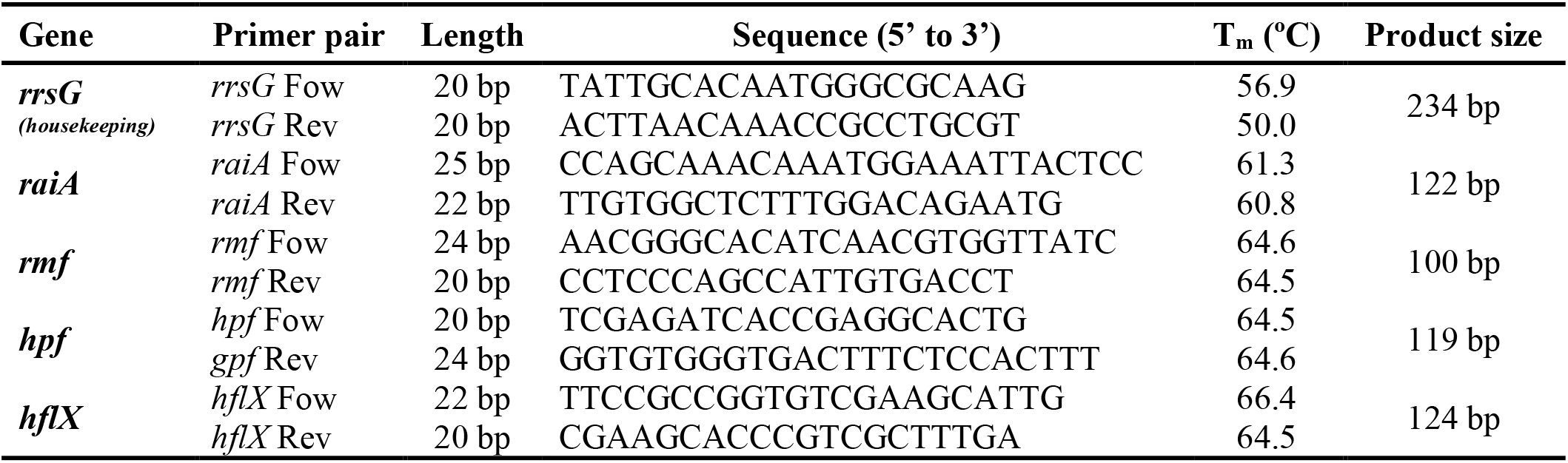
qRT-PCR primers.

**Table S3.** MqsR/MqsA/MqsC inhibits T2 phage by forming persister cells. Cells were contacted first with T2 phage at 0.1 MOI for 1 h followed by treatment with antibiotics: 10 MIC (100 µg/ml) for ampicillin, 100 MIC for ciprofloxacin (5 µg/ml), and 5 MIC for mitomycin C (10 µg/ml) for 3 h. Bold results indicate million-fold difference for inhibiting T2 phage. One average deviation shown. Survival percentages are normalized based on whether phage was used as indicated so either based on (i) no phage addition after 1 hour or based on (ii) adding phage after 1 hour. Note the presence of pCV1 and pCV1-*mqsRAC* increase ampicillin tolerance over 100-fold compared to the host MG1655, so ampicillin tolerance is greater than expected in the absence of phage.

| Strain | Elapsed time, h | Condition | CFU/ml average | Survival, % |
| --- | --- | --- | --- | --- |
| MG1655/pCV1 | 0 | Starting CFU | $1.4 \pm 0.2 \times 10^8$ | |
| | 1 | No phage | $1.8 \pm 0.5 \times 10^9$ | 100% |
|  | 1 | Phage T2 | **$3.4 \pm 0.8 \times 10^2$** | 100% |
| | 4 | No phage + ampicillin | $1 \pm 1 \times 10^8$ | $7 \pm 3\%$ |
| | 4 | No phage + ciprofloxacin | $2.4 \pm 1 \times 10^6$ | $0.20 \pm 0.08\%$ |
| | 4 | Phage T2 + ampicillin | 0.00 | $0 \pm 0\%$ |
| | 4 | Phage T2 + ciprofloxacin | 0.00 | $0 \pm 0\%$ |
| MG1655/pCV1-*mqsRAC* | 0 | Starting CFU | $1.9 \pm 0.3 \times 10^8$ | |
| | 1 | No phage | $1.5 \pm 0.2 \times 10^9$ | 100% |
|  | 1 | Phage T2 | **$4 \pm 1 \times 10^8$** | 100% |
| | 4 | No phage + ampicillin | $2.5 \pm 0.7 \times 10^8$ | $16 \pm 6\%$ |
| | 4 | No phage + ciprofloxacin | $3 \pm 1 \times 10^6$ | $0.20 \pm 0.09\%$ |
| | 4 | No phage + mitomycin C | $8 \pm 9 \times 10^1$ | $0.000005 \pm 0\%$ |
| | 4 | Phage T2 + ampicillin | $4 \pm 1 \times 10^7$ | $8 \pm 3\%$ |
| | 4 | Phage T2 + ciprofloxacin | $1.0 \pm 0.4 \times 10^6$ | $0.3 \pm 0.1\%$ |
| | 4 | Phage T2 + mitomycin C | $2 \pm 1 \times 10^2$ | $0.000034 \pm 0\%$ |

**Table S4.** Efficiency of plating and efficiency (EOP) of the center of infection assay (ECOI). (**A**) EOP assay with the *E. coli* strains used in this manuscript. (**B**) ECOI assay with the strains used in this manuscript. The efficiencies are relative to the strain marked ‘host’. One average deviation shown.

**A.**
| Strain | PFU/mL | EOP vs host | EOP vs empty plasmid |
| --- | --- | --- | --- |
| MG1655 (host) | $9 \pm 2 \times 10^9$ | 1 | |
| BW25113/pCV1 | $5.2 \pm 0.2 \times 10^9$ | $6.0 \times 10^{-1}$ | 1 |
| BW25113/pCV1-*mqsRAC* | $1.9 \pm 0.3 \times 10^6$ | $2.2 \times 10^{-4}$ | $3.7 \times 10^{-4}$ |
| MG1655/pCV1 | $3.7 \pm 0.4 \times 10^9$ | $4.3 \times 10^{-1}$ | 1 |
| MG1655/pCV1-*mqsRAC* | $1.30 \pm 0.07 \times 10^6$ | $1.5 \times 10^{-4}$ | $3.5 \times 10^{-4}$ |
| BW25113 $\Delta mcrB$/pCV1 | $4.0 \pm 0.5 \times 10^{10}$ | $4.6 \times 10^{-0}$ | 1 |
| BW25113 $\Delta mcrB$/pCV1-*mqsRAC* | $1.6 \pm 3 \times 10^8$ | $1.8 \times 10^{-2}$ | $4.0 \times 10^{-3}$ |

**B.**
| Strain | PFU/mL | ECOI vs host | EOP vs empty plasmid |
| --- | --- | --- | --- |
| MG1655 (host) | $3.7 \pm 0.1 \times 10^7$ | 1.000 | |
| BW25113/pCV1 | $1.43 \pm 0.07 \times 10^7$ | 0.387 | 1.000 |
| BW25113/pCV1-*mqsRAC* | $2.9 \pm 0.5 \times 10^6$ | 0.078 | 0.201 |
| MG1655/pCV1 | $2.6 \pm 0.4 \times 10^7$ | 0.691 | 1.000 |
| MG1655/pCV1-*mqsRAC* | $1.8 \pm 0.8 \times 10^6$ | 0.048 | 0.069 |
| BW25113 $\Delta mcrB$/pCV1 | $2 \pm 1 \times 10^8$ | 5.495 | 1.000 |
| BW25113 $\Delta mcrB$/pCV1-*mqsRAC* | $9 \pm 7 \times 10^6$ | 0.252 | 0.046 |

**Table S5.** Percentage of *E. coli* persister cell resuscitation after treatment with T2 phage (0.1 MOI) for 1 h. Values in bold are included in the main text, and values for the number of cells with the indicated phenotype are listed parenthetically.

| Sample | No of cells | % of waking in 0.5 h | % of waking in 0.5 to 3 h | % Elongated cells | % of dead |
| --- | --- | --- | --- | --- | --- |
| 1 | 33 | - | 88% (29/33) | 9% (3/33) | 3 % (1/33) |
| 2 | 25 | 20% (5/25) | 12% (3/25) | 16% (4/25) | 52 % (13/25) |
| **3** | **27** | **41% (11/27)** | **33% (9/27)** | **11% (3/27)** | **15 % (4/27)** |
| 4 | 66 | 58% (38/66) | 15% (10/66) | 18% (12/66) | 9 % (6/66) |
| 5 | 24 | 25% (6/24) | 21% (5/24) | 33% (8/24) | 21 % (5/24) |
| **Average** |  | 36 ± 13% | 34 ± 22% | 18 ± 7% | 20 ± 13% |

**Table S6.** Phage attack does not induce ribosome inactivation proteins. Fold changes for the wild-type *E. coli* with the empty plasmid pCV1 relative to the wild-type producing MqsR/MqsA/MqsC (from pCV1-*mqsRAC*) after 15 min of T2 attack as determined by qRT-PCR. Cycle numbers (C_t_) are indicated for each sample including that for the target genes as well as that of the house-keeping gene, *rrsG,* which was used to normalize the data. Fold were calculated as described earlier (58): 2^-((C_t *pCV1,i*_ *-* C_t *rrsG-CV1,i*_) - (C_t *mqsRAC,i*_ - C_t *rrsG-mqsRAC,i*_)) where C_t_ *_mqsRAC,i_*___is the cycle number for *rmf, hpf,* or *raiA* mRNA in MG1655*/*pCV1-mqsRAC, C_t_ *_pCV1,i_*___is the cycle number for *rmf, hpf,* or *raiA* mRNA in MG1655*/*pCV1, C_t_ *_rrsG-mqsRAC,i_* is the cycle number for the associated housekeeping gene *rrsG* for *rmf, hpf,* or *raiA* mRNA in MG1655*/pCV1-mqsRAC*, C_t_ *_rrsG-pCV1i_* is cycle number for the associated housekeeping gene *rrsG* for *rmf, hpf,* or *raiA* mRNA in MG1655*/pCV1*, ΔC_T_ is the difference of the target gene cycle number and the housekeeping cycle number, ΔΔC_T_ is the difference of the MG1655*/pCV1-mqsRAC* ΔC_T_ vs. the MG1655*/pCV1* ΔC_T_, and fold indicates the fold change of the relative expression of the target gene in MG1655*/pCV1-mqsRAC* vs. MG1655*/pCV1*.

| Gene | *rrsG* |  | *rmf* |  | *hpf* |  | *raiA* |  |
| --- | --- | --- | --- | --- | --- | --- | --- | --- |
| Strain | MG1655<br>/pCV1 | MG1655<br>/pCV1-<br>*mqsRAC* | MG1655<br>/pCV1 | MG1655<br>/pCV1-<br>*mqsRAC* | MG1655<br>/pCV1 | MG1655<br>/pCV1-<br>*mqsRAC* | MG1655<br>/pCV1 | MG1655<br>/pCV1-<br>*mqsRAC* |
| $C_T$ | 4.6<br>$\pm 0.5$ | 4.48<br>$\pm 0.05$ | 16.9<br>$\pm 0.6$ | 17.0<br>$\pm 0.2$ | 15.3<br>$\pm 0.6$ | 15.2<br>$\pm 0.1$ | 12.8<br>$\pm 0.2$ | 14.7<br>$\pm 0.1$ |
| $\Delta C_T$ | | | 12.3<br>$\pm 0.1$ | 12.6<br>$\pm 0.2$ | 10.7<br>$\pm 0.2$ | 10.7<br>$\pm 0.1$ | 8.2<br>$\pm 0.3$ | 10.2<br>$\pm 0.2$ |
| $\Delta\Delta C_T$ | | | 0.3<br>$\pm 0.1$ | | 0.02<br>$\pm 0.02$ | | 2.0<br>$\pm 0.1$ | |
| Fold |  |  | 1.2 |  | 1.05 |  | 4.0 |  |

**Table S7.** MqsR/MqsA/MqsC inhibits T2 phage through cooperation with restriction/modification systems. (**A**) Survival after 30 min of T2 phage infection at 0.1 MOI. (**B**) BW25113 and BW25113 *mcrB* cells producing MqsR/MqsA/MqsC from its native promoter (and the empty plasmid negative control) were contacted first with T2 phage at 0.1 MOI for 1 h followed by treatment with ampicillin (10 MIC, 100 µg/ml) or mitomycin C (5 MIC, 10 µg/ml) for 3 h. Bold indicates CFU comparison of persister cells. Survival percentages are normalized based on whether phage was used as indicated so either based on (i) no phage addition after 1 hour or based on (ii) adding phage after 1 hour. Note the presence of pCV1 and pCV1-*mqsRAC* increase ampicillin tolerance over 100-fold compared to the host MG1655, so ampicillin tolerance is greater than expected in the absence of phage. (**C**) Phage titers after 30 min of infection at 0.1 MOI. One average deviation shown. Part **A** includes results from two independent cultures whereas parts **B** and **C** include results from four independent cultures each.

| A | Strain | CFU/ml average |
| --- | --- | --- |
| | BW25113 | $1.4 \pm 0.3 \times 10^6$ |
| | BW25113 $\Delta mrr$ | $8 \pm 2 \times 10^4$ |
| | BW25113 $\Delta recD$ | $4.0 \pm 0.4 \times 10^6$ |
| | BW25113 $\Delta mcrB$ | $2 \pm 1 \times 10^2$ |

| B | Strain | Condition | CFU/ml average | Survival % |
| --- | --- | --- | --- | --- |
| | BW25113/pCV1 | Starting CFU | $1.2 \pm 0.2 \times 10^8$ | |
| | | No phage | $1.6 \pm 0.1 \times 10^9$ | 100% |
|  |  | Phage T2 | **$3 \pm 1 \times 10^2$** | 100% |
| | | No phage + ampicillin | $2.5 \pm 0.5 \times 10^8$ | $15 \pm 6\%$ |
| | | Phage T2 + ampicillin | 0.00 | $0 \pm 0\%$ |
| | BW25113 $\Delta mcrB$/pCV1 | Starting CFU | $1.5 \pm 0.3 \times 10^8$ | |
| | | No phage | $1.7 \pm 0.3 \times 10^9$ | 100% |
|  |  | Phage T2 | **$6 \pm 6 \times 10^1$** | 100% |
| | | No phage + ampicillin | $1.7 \pm 0.3 \times 10^7$ | $8 \pm 4\%$ |
| | | Phage T2 + ampicillin | 0.00 | $0 \pm 0\%$ |
| | BW25113/pCV1-*mqsRAC* | Starting CFU | $1.2 \pm 0.2 \times 10^8$ | |
| | | No phage | $1.7 \pm 0.3 \times 10^9$ | 100% |
| | | Phage T2 | $3.8 \pm 0.3 \times 10^8$ | 100% |
| | | No phage + ampicillin | $2.0 \pm 0.3 \times 10^8$ | $12 \pm 3\%$ |
| | | No phage + mitomycin C | $4 \pm 3 \times 10^2$ | $0.000032 \pm 0\%$ |
| | | Phage T2 + ampicillin | **$1.7 \pm 0.5 \times 10^6$** | $0.4 \pm 0.1\%$ |
| | | Phage T2 + mitomycin C | $1.4 \pm 1 \times 10^2$ | $0.00010 \pm 0\%$ |
| | BW25113 $\Delta mcrB$/pCV1-*mqsRAC* | Starting CFU | $1.4 \pm 0.3 \times 10^8$ | |
| | | No phage | $8 \pm 2 \times 10^8$ | 100% |
| | | Phage T2 | $1.3 \pm 0.6 \times 10^7$ | 100% |
| | | No phage + ampicillin | $1.9 \pm 0.5 \times 10^7$ | $3 \pm 1\%$ |
| | | No phage + mitomycin C | $1.1 \pm 0.9 \times 10^3$ | $0.00012 \pm 0\%$ |
| | | Phage T2 + ampicillin | **$2 \pm 2 \times 10^5$** | $1.6 \pm 0.7\%$ |
| | | Phage T2 + mitomycin C | $4 \pm 5 \times 10^1$ | $0.000078 \pm 0\%$ |

| C | Strain | PFU/ml average | CFU/ml average |
| --- | --- | --- | --- |
| | BW25113/pCV1-*mqsRAC* | $7 \pm 2 \times 10^5$ | $1.8 \pm 0.7 \times 10^8$ |
| | BW25113 $\Delta mcrB$/pCV1-*mqsRAC* | $2.9 \pm 0.9 \times 10^6$ | $3 \pm 1 \times 10^7$ |

**Supplemental Figure S1.**
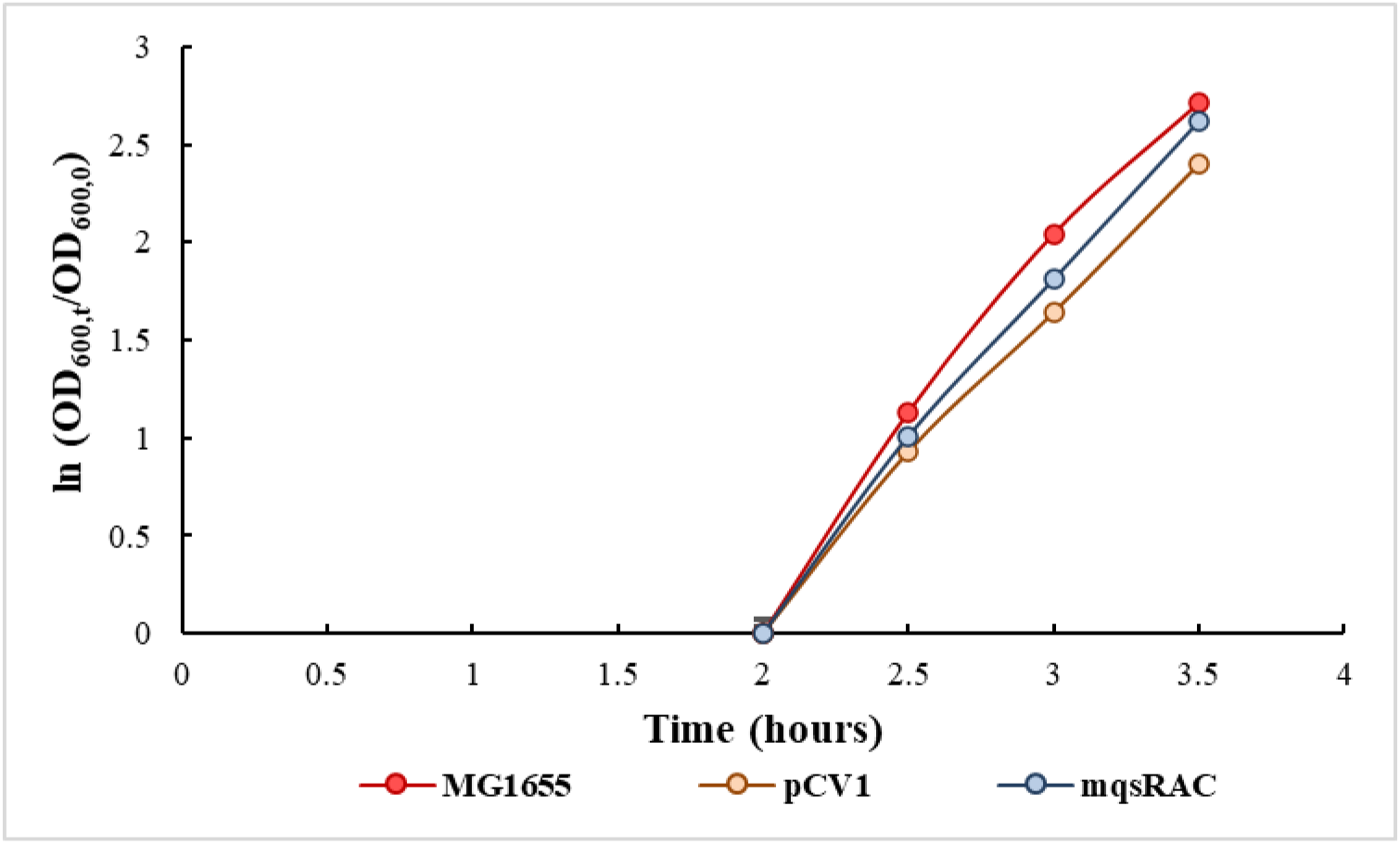
Growth of four independent colonies of each strain in LB at 37°C. MG1655, the wild-type host, is indicated in red, MG1655/pCV1, the host with the empty plasmid (negative control), is indicated in orange, and MG1655/pCV1-mqsRAC, the host with the MqsR/MqsA/MqsC tripartite toxin/antitoxin system, is indicated in blue. One average deviation shown.

**Supplemental Figure S2.**
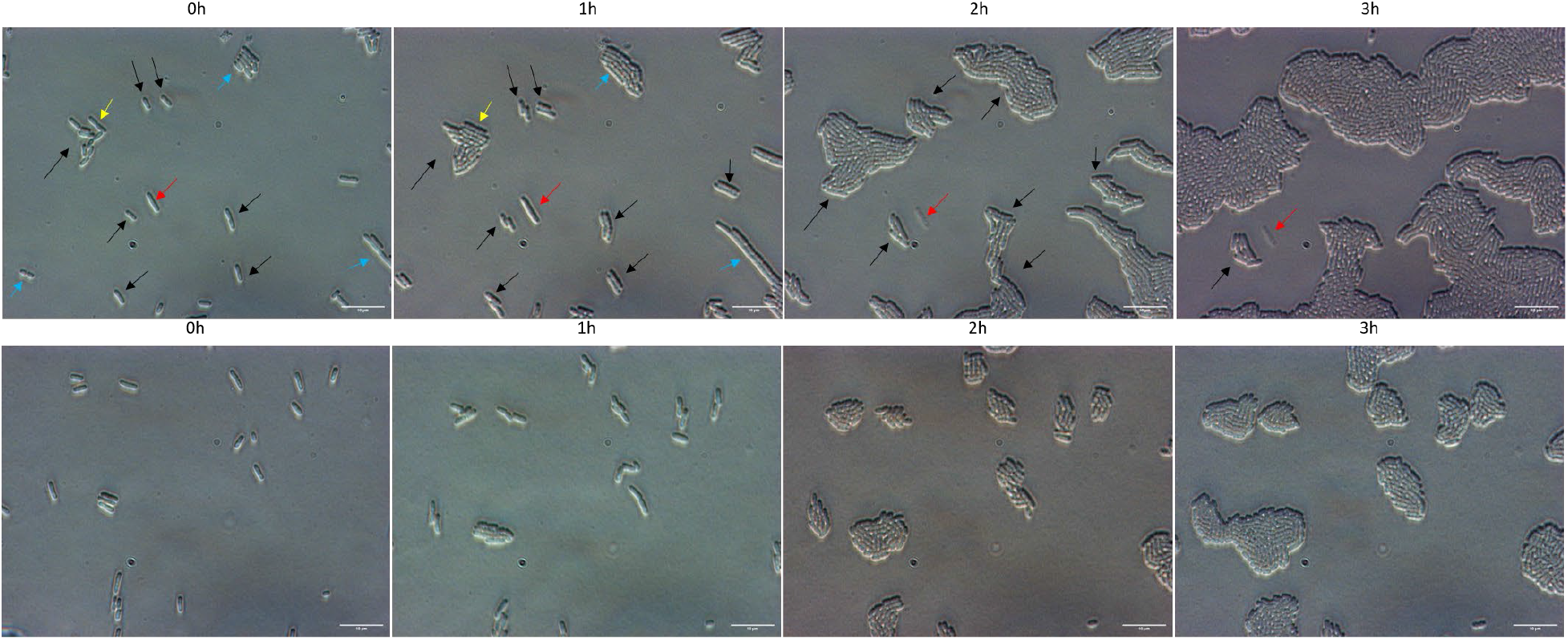
Long-term heterogenous single-cell resuscitation after phage attack. Representative images (from five independent cultures) of the resuscitation of *E. coli* persister cells with MqsR/MqsA/MqsC (**upper row**) and division of exponential cells (**lower row**) after 0 to 3 h as determined with light microscopy (Zeiss Axio Scope.A1) using LB agarose gel pads. The persister cells were generated by the addition of T2 phage at 0.1 MOI for 1 hour. Cells with the empty plasmid (i.e., no MqsR/MqsA/MqsC) are not shown due to the cellular debris that stems from complete eradication by T2 phage. Black arrows indicate cells with immediate waking (within 30 min), yellow arrows indicate cells with delayed waking (waking between 30 – 180 min), red arrows indicate cells that wake then die (lyse within 3 h), and blue arrows indicate cells that elongate. Black arrows were not added to exponential cultures pictures because all of them divide. Data for percentages from 0 to 1 h are shown in **Table S5**.

**Supplemental Figure S3.**
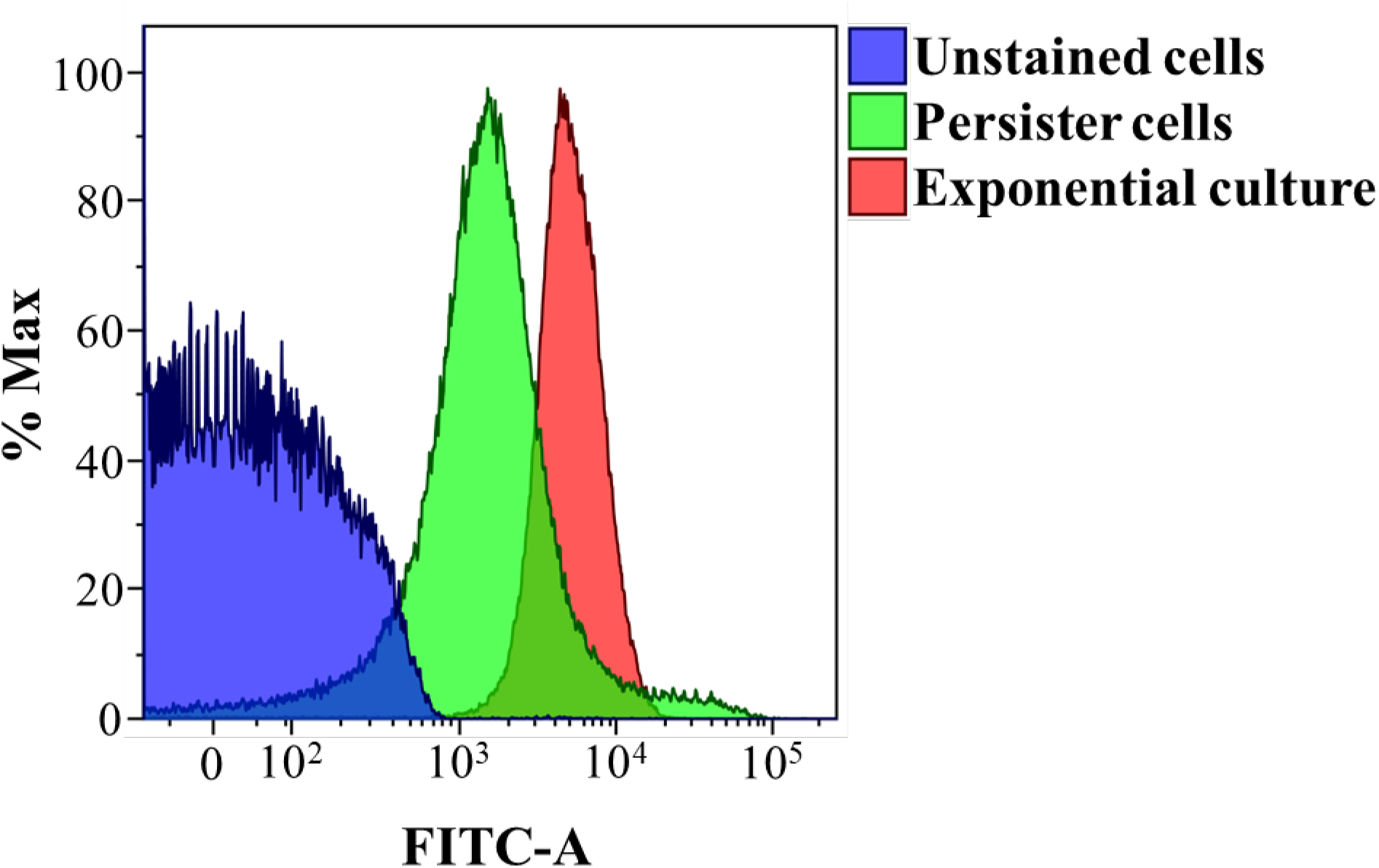
Metabolic activity via flow cytometry. MG1655/pCV1*-mqsRAC* was grown to a turbidity of 0.5 at 600 nm and T2 phage was added (MOI 0.1) for 1 h, then stained for metabolic activity using the BacLight™ RedoxSensor™ Green Vitality Kit (‘persister’ cells, green) and compared to exponentially-growing cells stained with the same kit (‘Exponential culture’, red). RedoxSensor Green measures the cellular redox state. ‘Unstained’ indicates results with an overnight, unstained sample. ‘%Max’ indicates the relative number of cells (100,000 cells total), and ‘FITC-A’ is indicative of the intensity of the RedoxSensor Green fluorescence. One representative image of two independent cultures shown.

**Supplemental Figure S4.**
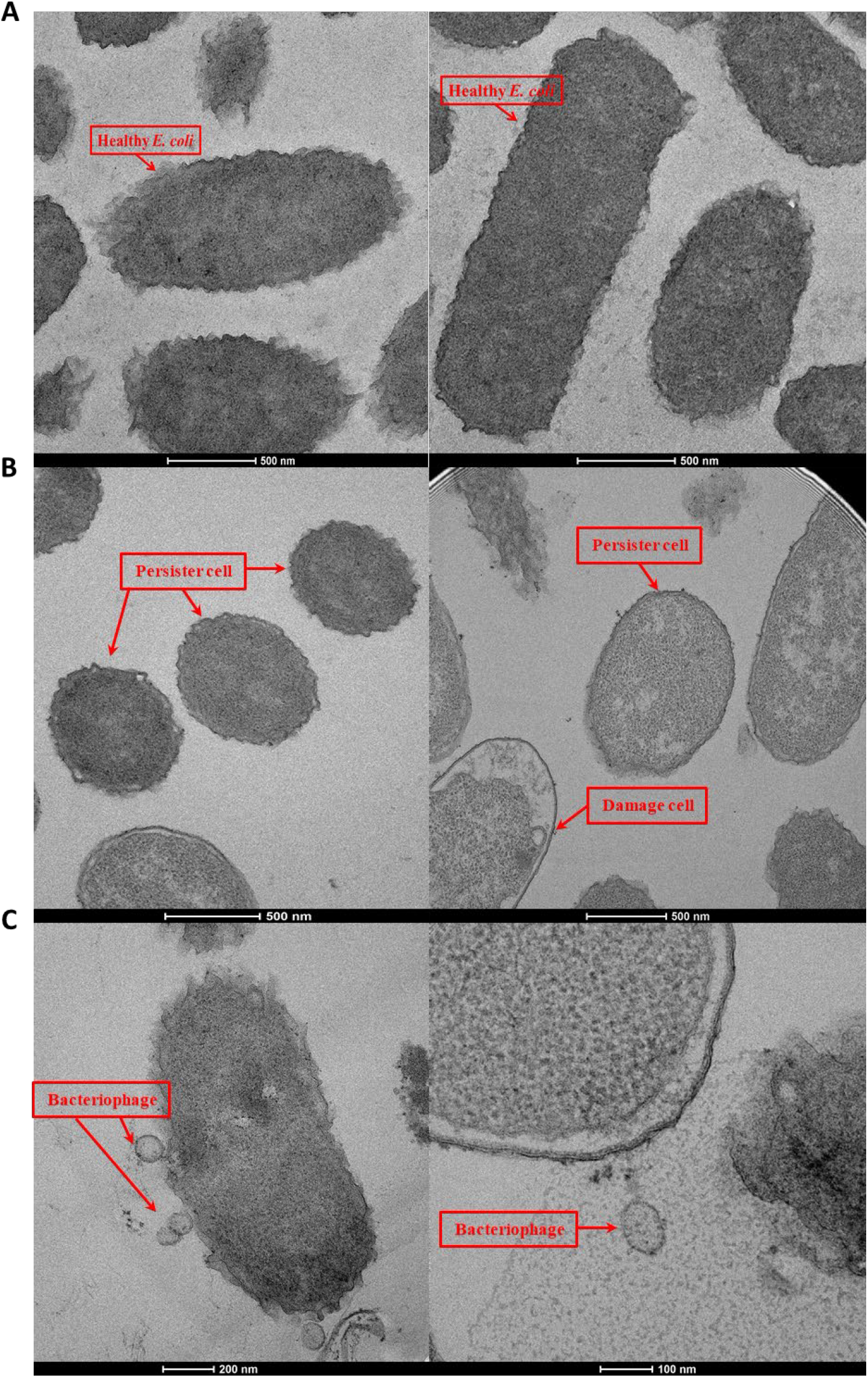
Transmission electron microscopy of *E. coli* cells after T2 phage attack. MG1655/pCV1-mqsRAC was grown to a turbidity of 0.5 at 600 nm, and T2 phage was added (MOI 0.1) for 1 h. (**A**) exponentially-growing cells prior to phage addition, (**B**) persister cells formed after T2 attack, which are smaller, and spheroid compared to exponentially-growing cells, and (**C**) damaged cells after T2 phage infection. Phage T2 indicated with an arrow and labelled ‘Bacteriophage’.

## REFERENCES

1. Keen, E.C. (2015) A century of phage research: Bacteriophages and the shaping of modern biology. BioEssays, 37, 6–9.

2. van Beljouw, S.P.B., Sanders, J., Rodríguez-Molina, A. and Brouns, S.J.J. (2022) RNA-targeting CRISPR–Cas systems. Nature Rev Microbiol.

3. Meeske, A.J., Nakandakari-Higa, S. and Marraffini, L.A. (2019) Cas13-induced cellular dormancy prevents the rise of CRISPR-resistant bacteriophage. Nature, 570, 241–245.

4. Dimitriu, T., Kurilovich, E., Łapińska, U., Severinov, K., Pagliara, S., Szczelkun, M.D. and Westra, E.R. (2022) Bacteriostatic antibiotics promote CRISPR-Cas adaptive immunity by enabling increased spacer acquisition. Cell Host Microbe, 30, 31–40.e35.

5. Hobby, G.L., Meyer, K. and Chaffee, E. (1942) Observations on the Mechanism of Action of Penicillin. Exp Biol Med 50, 281–285.

6. Kim, J.-S., Chowdhury, N., Yamasaki, R. and Wood, T.K. (2018) Viable But Non-Culturable and Persistence Describe the Same Bacterial Stress State. Environ Microbiol, 20, 2038–2048.

7. Hong, S.H., Wang, X., O’Connor, H.F., Benedik, M.J. and Wood, T.K. (2012) Bacterial persistence increases as environmental fitness decreases. Microbial Biotechnol, 5, 509–522.

8. Korch, S.B., Henderson, T.A. and Hill, T.M. (2003) Characterization of the *hipA7* allele of *Escherichia coli* and evidence that high persistence is governed by (p)ppGpp synthesis. Mol Microbiol, 50, 1199–1213.

9. Song, S. and Wood, T.K. (2020) ppGpp Ribosome Dimerization Model for Bacterial Persister Formation and Resuscitation. Biochem Biophys Res Com, 523, 281–286.

10. Yamasaki, R., Song, S., Benedik, M.J. and Wood, T.K. (2020) Persister Cells Resuscitate Using Membrane Sensors that Activate Chemotaxis, Lower cAMP Levels, and Revive Ribosomes. iScience, 23, 100792.

11. Kwan, B.W., Valenta, J.A., Benedik, M.J. and Wood, T.K. (2013) Arrested protein synthesis increases persister-like cell formation. Antimicrob Agents Chemother, 57, 1468–1473.

12. Chowdhury, N., Kwan, B.W. and Wood, T.K. (2016) Persistence Increases in the Absence of the Alarmone Guanosine Tetraphosphate by Reducing Cell Growth. Sci Rep, 6, 20519.

13. Wang, X., Lord, D.M., Cheng, H.-Y., Osbourne, D.O., Hong, S.H., Sanchez-Torres, V., Quiroga, C., Zheng, K., Herrmann, T., Peti, W. et al. (2012) A Novel Type V TA System Where mRNA for Toxin GhoT is Cleaved by Antitoxin GhoS. Nature Chem Biol, 8, 855–861.

14. Pecota, D.C. and Wood, T.K. (1996) Exclusion of T4 Phage by the *hok/sok* Killer Locus from Plasmid R1. J Bacteriol, 178, 2044–2050.

15. Lau, R.K., Enustun, E., Gu, Y., Nguyen, J.V. and Corbett, K.D. (2022) A conserved signaling pathway activates bacterial CBASS immune signaling in response to DNA damage. EMBO J, e111540.

16. Duncan-Lowey, B., Tal, N., Johnson, A.G., Rawson, S., L.Mayer, M., Doron, S., Millman, A., Melamed, S., Fedorenko, T., Kacen, A., et al. (2023) Cryo-EM structure of the RADAR supramolecular anti-phage defense complex. Cell, 186, 987–998.e15

17. Leavitt, A., Yirmiya, E., Amitai, G., Lu, A., Garb, J., Herbst, E., Morehouse, B.R., Hobbs, S.J., Antine, S.P., Sun, Z.-Y.J. et al. (2022) Viruses inhibit TIR gcADPR signaling to overcome bacterial defense. Nature, 611, 326–331.

18. Bordes, P., Sala, A.J., Ayala, S., Texier, P., Slama, N., Cirinesi, A.-M., Guillet, V., Mourey, L. and Genevaux, P. (2016) Chaperone addiction of toxin–antitoxin systems. Nature Comm, 7, 13339.

19. Vassallo, C.N., Doering, C.R., Littlehale, M.L., Teodoro, G.I.C. and Laub, M.T. (2022) A functional selection reveals previously undetected anti-phage defence systems in the *E. coli* pangenome. Nature Microbiol, 7, 1568–1579.

20. Brown, B.L., Grigoriu, S., Kim, Y., Arruda, J.M., Davenport, A., Wood, T.K., Peti, W. and Page, R. (2009) Three dimensional structure of the MqsR:MqsA complex: A novel toxin:antitoxin pair comprised of a toxin homologous to RelE and an antitoxin with unique properties. PLoS Pathog, 5, e1000706.

21. Ren, D., Bedzyk, L.A., Thomas, S.M., Ye, R.W. and Wood, T.K. (2004) Gene expression in *Escherichia coli* biofilms. Appl Microbiol Biotechnol, 64, 515–524.

22. Kwan, B.W., Chowdhury, N. and Wood, T.K. (2015) Combatting Bacterial Infections by Killing Persister Cells with Mitomycin C. Environ Microbiol, 17 4406–4414.

23. Kim, J.-S., Yamasaki, R., Song, S., Zhang, W. and Wood, T.K. (2018) Single Cell Observations Show Persister Cells Wake Based on Ribosome Content. Environ Microbiol, 20, 2085–2098.

24. Kutter, E. (2009) In Martha R. J. Clokie, A. M. K. (ed.), Bacteriophages: Methods and Protocols, Volume 1: Isolation, Characterization, and Interactions. Humana Press, a part of Springer Science+Business Media, Vol. 501.

25. Sing, W.D. and Klaenhammer, T.R. (1990) Characteristics of phage abortion conferred in lactococci by the conjugal plasmid pTR2030. Microbiol, 136, 1807–1815.

26. Pacios, O., Fernández-García, L., Bleriot, I., Blasco, L., Ambroa, A., López, M., Ortiz-Cartagena, C., Cuenca, F.F., Oteo-Iglesias, J., Pascual, Á. et al. (2021) Phenotypic and Genomic Comparison of Klebsiella pneumoniae Lytic Phages: vB_KpnM-VAC66 and vB_KpnM-VAC13. Viruses, 14.

27. Bleriot, I., Blasco, L., Pacios, O., Fernández-García, L., López, M., Ortiz-Cartagena, C., Barrio-Pujante, A., Fernández-Cuenca, F., Pascual, Á., Martínez-Martínez, L. et al. (2023) Proteomic Study of the Interactions between Phages and the Bacterial Host Klebsiella pneumoniae. Microbiol Spectr, 11, e0397422.

28. Song, S., Gong, T., Yamasaki, R., Kim, J.S. and Wood, T.K. (2019) Identification of a potent indigoid persister antimicrobial by screening dormant cells. Biotechnol Bioeng, 116, 2263–2274.

29. Song, S., Kim, J.-S., Yamasaki, R., Oh, S., Benedik, M.J. and Wood, T.K. (2021) *Escherichia coli* cryptic prophages sense nutrients to influence persister cell resuscitation. Environ Microbiol, 23, 7245–7254.

30. Song, S. and Wood, T.K. (2020) Persister Cells Resuscitate via Ribosome Modification by 23S rRNA Pseudouridine Synthase RluD. Environ Microbiol, 22, 850–857.

31. Wood, T.K., Knabel, S.J. and Kwan, B.W. (2013) Bacterial persister cell formation and dormancy. Appl Environ Microbiol, 79, 7116–7121.

32. Shan, Y., Brown Gandt, A., Rowe, S.E., Deisinger, J.P., Conlon, B.P. and Lewis, K. (2017) ATP-Dependent Persister Formation in *Escherichia coli*. mBio, 8.

33. Sulaiman, J.E. and Lam, H. (2020) Proteomic Investigation of Tolerant Escherichia coli Populations from Cyclic Antibiotic Treatment. J Proteome Res, 19, 900–913.

34. Lewis, K. (2007) Persister cells, dormancy, and infectious disease. Nature Rev Microbiol, 5, 48–56.

35. Lewis, K. (2008) Multidrug tolerance of biofilms and persister cells. Curr Top Microbiol Immunol, 322, 107–131.

36. Bigger, J.W. (1944) Treatment of Staphylococcal Infections with Penicillin by Intermittent Sterilisation. Lancet, 244, 497–500.

37. Pu, Y., Li, Y., Jin, X., Tian, T., Ma, Q., Zhao, Z., Lin, S.-y., Chen, Z., Li, B., Yao, G., et al. (2019) ATP-Dependent Dynamic Protein Aggregation Regulates Bacterial Dormancy Depth Critical for Antibiotic Tolerance. Mol Cell, 73, 143–156.

38. Goormaghtigh, F. and Van Melderen, L. (2019) Single-cell imaging and characterization of *Escherichia coli* persister cells to ofloxacin in exponential cultures. Science Adv, 5, eaav9462.

39. Şimşek, E. and Kim, M. (2019) Power-law tail in lag time distribution underlies bacterial persistence. PNAS, 116, 17635–17640.

40. Windels, E.M., Ben Meriem, Z., Zahir, T., Verstrepen, K.J., Hersen, P., Van den Bergh, B. and Michiels, J. (2019) Enrichment of persisters enabled by a ß-lactam-induced filamentation method reveals their stochastic single-cell awakening. Comm Biol, 2, 426.

41. Levinthal, C. and Visconti, N. (1953) Growth and Recombination in Bacterial Viruses. Genetics, 38, 500–511.

42. Little, J.S., Dedrick, R.M., Freeman, K.G., Cristinziano, M., Smith, B.E., Benson, C.A., Jhaveri, T.A., Baden, L.R., Solomon, D.A. and Hatfull, G.F. (2022) Bacteriophage treatment of disseminated cutaneous Mycobacterium chelonae infection. Nature Comm, 13, 2313.

43. Cruz-Muñiz, M.Y., López-Jacome, L.E., Hernández-Durán, M., Franco-Cendejas, R., Licona-Limón, P., Ramos-Balderas, J.L., Martinéz-Vázquez, M., Belmon-Díazt, J.A., Wood, T.K. and García-Contreras, R. (2017) Repurposing the anticancer drug mitomycin C for the treatment of persistent *Acinetobacter baumannii* infections. Int J Antimicrob Agents, 49, 88–92.

44. Williams, M.C., Reker, A.E., Margolis, S.R., Liao, J., Wiedmann, M., Rojas, E.R. and Meeske, A.J. (2023) Restriction endonuclease cleavage of phage DNA enables resuscitation from Cas13-induced bacterial dormancy. Nature Microbiol, 8, 400–409.

45. Tesson, F., Hervé, A., Mordret, E., Touchon, M., d’Humières, C., Cury, J. and Bernheim, A. (2022) Systematic and quantitative view of the antiviral arsenal of prokaryotes. Nature Comm, 13, 2561.

46. Thompson, M.G., Sedaghatian, N., Barajas, J.F., Wehrs, M., Bailey, C.B., Kaplan, N., Hillson, N.J., Mukhopadhyay, A. and Keasling, J.D. (2018) Isolation and characterization of novel mutations in the pSC101 origin that increase copy number. Sci Reports, 8, 1590.

47. Lopatina, A., Tal, N. and Sorek, R. (2020) Abortive Infection: Bacterial Suicide as an Antiviral Immune Strategy. Ann Rev Virol, 7, 371–384.

48. LeRoux, M. and Laub, M.T. (2022) Toxin-Antitoxin Systems as Phage Defense Elements. Ann Rev Microbiol, 76, 21–43.

49. LeRoux, M., Srikant, S., Teodoro, G.I.C., Zhang, T., Littlehale, M.L., Doron, S., Badiee, M., Leung, A.K.L., Sorek, R. and Laub, M.T. (2022) The DarTG toxin-antitoxin system provides phage defence by ADP-ribosylating viral DNA. Nature Microbiol, 7, 1028–1040.

50. Arias, C.F., Acosta, F.J., Bertocchini, F., Herrero, M.A. and Fernández-Arias, C. (2022) The coordination of anti-phage immunity mechanisms in bacterial cells. Nature Commun, 13, 7412.

51. Rousset, F., Yirmiya, E., Nesher, S., Brandis, A., Mehlman, T., Itkin, M., Malitsky, S., Millman, A., Melamed, S. and Sorek, R. (2023) A conserved family of immune effectors cleaves cellular ATP upon viral infection. bioRxiv, 2023.2001.2024.525353.

52. Conlon, B.P., Rowe, S.E., Gandt, A.B., Nuxoll, A.S., Donegan, N.P., Zalis, E.A., Clair, G., Adkins, J.N., Cheung, A.L. and Lewis, K. (2016) Persister formation in *Staphylococcus aureus* is associated with ATP depletion. Nature Microbiol, 1, 16051.

53. Cheng, H.-Y., Soo, V.W.C., Islam, S., McAnulty, M.J., Benedik, M.J. and Wood, T.K. (2014) Toxin GhoT of the GhoT/GhoS toxin/antitoxin system damages the cell membrane to reduce adenosine triphosphate and to reduce growth under stress. Environ Microbiol, 16, 1741–1754.

54. Ho, P., Chen, Y., Biswas, S., Canfield, E. and Feldman, D.E. (2023) Bacteriophage anti-defense genes that neutralize TIR and STING immune responses. Cell Reports, 42, 112305.

55. Makarova, K.S., Anantharaman, V., Aravind, L. and Koonin, E.V. (2012) Live virus-free or die: coupling of antivirus immunity and programmed suicide or dormancy in prokaryotes. Biol Direct, 7, 40.

56. Blattner, F.R., III Plunkett, G., Bloch, C.A., Perna, N.T., Burland, V., Riley, M., Collado-Vides, J., Glasner, J.D., Rode, C.K., Mayhew, G.F., et al. (1997) The complete genome sequence of *Escherichia coli* K-12. Science, 277, 1453–1462.

57. Baba, T., Ara, T., Hasegawa, M., Takai, Y., Okumura, Y., Baba, M., Datsenko, K.A., Tomita, M., Wanner, B.L. and Mori, H. (2006) Construction of *Escherichia coli* K-12 in-frame, single-gene knockout mutants: the Keio collection. Mol Syst Biol, 2, 2006 0008.

58. Pfaffl, M.W. (2001) A new mathematical model for relative quantification in real-time RT-PCR. Nucleic Acids Res., 29, e45.

